# Cellular development and evolution of the mammalian cerebellum

**DOI:** 10.1101/2021.12.20.473443

**Authors:** Mari Sepp, Kevin Leiss, Ioannis Sarropoulos, Florent Murat, Konstantin Okonechnikov, Piyush Joshi, Evgeny Leushkin, Noe Mbengue, Céline Schneider, Julia Schmidt, Nils Trost, Lisa Spänig, Peter Giere, Philipp Khaitovich, Steven Lisgo, Miklós Palkovits, Lena M. Kutscher, Simon Anders, Margarida Cardoso-Moreira, Stefan M. Pfister, Henrik Kaessmann

**Affiliations:** Center for Molecular Biology of Heidelberg University (ZMBH), DKFZ-ZMBH Alliance, Heidelberg, Germany; Hopp-Children’s Cancer Center Heidelberg (KiTZ), Heidelberg, Germany; Division of Pediatric Neurooncology, German Cancer Research Center (DKFZ) and German Cancer Consortium (DKTK), Heidelberg, Germany; Developmental Origins of Pediatric Cancer Junior Group, German Cancer Research Center (DKFZ) and German Cancer Consortium (DKTK), Heidelberg, Germany; Museum für Naturkunde Berlin, Leibniz Institute for Research on Evolution and Biodiversity Science at the Humboldt University Berlin, Berlin, Germany; Center for Neurobiology and Brain Restoration, Skolkovo Institute of Science and Technology, Moscow, Russia; Biosciences Institute, Newcastle University, Newcastle, UK; Human Brain Tissue Bank, Semmelweis University, Budapest, Hungary; BioQuant, Heidelberg University, Heidelberg, Germany; Evolutionary Developmental Biology Laboratory, Francis Crick Institute, London NW1 1AT, UK; Department of Pediatric Hematology and Oncology, Heidelberg University Hospital, Heidelberg, Germany

## Abstract

The expansion of the neocortex, one of the hallmarks of mammalian evolution^1,2^, was accompanied by an increase in the number of cerebellar neurons^3^. However, little is known about the evolution of the cellular programs underlying cerebellum development in mammals. In this study, we generated single-nucleus RNA-sequencing data for ∼400,000 cells to trace the development of the cerebellum from early neurogenesis to adulthood in human, mouse, and the marsupial opossum. Our cross-species analyses revealed that the cellular composition and differentiation dynamics throughout cerebellum development are largely conserved, except for human Purkinje cells. Global transcriptome profiles, conserved cell state markers, and gene expression trajectories across neuronal differentiation show that the cerebellar cell type-defining programs have been overall preserved for at least 160 million years. However, we also discovered differences. We identified 3,586 genes that either gained or lost expression in cerebellar cells in one of the species, and 541 genes that evolved new expression trajectories during neuronal differentiation. The potential functional relevance of these cross-species differences is highlighted by the diverged expression patterns of several human disease-associated genes. Altogether, our study reveals shared and lineage-specific programs governing the cellular development of the mammalian cerebellum, and expands our understanding of the evolution of mammalian organ development.

## MAIN TEXT

Establishing causal relationships between the molecular and phenotypic evolution of the nervous systems of our species and other mammals is a primary goal in biology. The expansion of the neocortex, considered as one of the hallmarks of mammalian evolution^1,2^, was accompanied by an increase in the number of cerebellar neurons^3^. The cerebellum varies substantially in size and shape across vertebrates^4^. In mammals, it contains more than half of the neurons of the entire brain^3^ and is involved in cognitive, affective, and linguistic processing besides its well-established role in sensory-motor control^5^. The cellular architecture of the adult cerebellum has long been viewed as being relatively simple, with its characteristic Purkinje and granule cells organized into cortical layers and the deep nuclei neurons embedded inside the white matter, but it is increasingly recognized to exhibit rather complex regional specializations^6–8^. Our understanding of mammalian cerebellum development stems mostly from rodent-based studies^6^, although differences in the cellular heterogeneity in the human cerebellum have been recognized^8,9^. Recent single-cell transcriptome studies of the developing mouse^10–12^ and human^13^ cerebellum provided new insights into gene expression programs in differentiating cerebellar cells, but an evolutionary analysis of the molecular and cellular diversity of the mammalian cerebellum across development is missing. In this study, we used single-nucleus RNA-sequencing (snRNA-seq) to study the development of the cerebellum from early neurogenesis to adulthood in three therian species: two eutherians (human, mouse) and a marsupial (opossum). Our analyses of these data, which provide an extensive resource (https://apps.kaessmannlab.org/sc-cerebellum-transcriptome), unveiled ancestral as well as species-specific cellular and molecular features of cerebellum development spanning ∼160 million years of mammalian evolution.

### Atlases of cerebellum development across mammals

The cerebellum has a protracted course of development, extending from early embryonic stages well into postnatal life^6^. To characterize the entire development of the mammalian cerebellum, we produced droplet-based snRNA-seq data for cerebella from 9-12 developmental stages in mouse, human, and opossum (Fig. 1a, Extended Data Fig.1a). We acquired high quality transcriptional profiles of 395,736 cells sequenced in 87 libraries, with a median of 21,181 uniquely mapping reads per library and 2,354 RNA molecules detected per cell (Extended Data Fig. 1a-d, Supplementary Table 1). We used LIGER^14^ to correct for batch effects and merge datasets from all stages for each species by integrative non-negative matrix factorization. We identified clusters by the Louvain algorithm and projected cells into a low-dimensional embedding (Extended Data Fig.1e). Because cerebellum development is best understood in mouse, we used known cell type markers^6,12^ and public *in situ* hybridisation data^15,16^ to build a hierarchical annotation of the mouse dataset, which we then transferred to the human and opossum datasets by the pairwise integration of the datasets within the orthologous gene expression space, and manual curation to account for biological and technical variance between the datasets (Extended Data Fig. 2a,b, Supplementary Tables 2-4). We grouped the cells into broad lineages based on their developmental origin, into cell types, and into cell differentiation states (hereafter referred to as cell states), consistent with the ongoing efforts in establishing cell ontologies^17^. Within cell types, we defined subtypes only at cell states that displayed remaining variability. Across the three species, we identified 25 cell types divided into 44 cell states, and for 12 cell states, we further split the cells into 49 subtypes (Fig. 1b-c, Extended Data Fig. 2c).

**Fig. 1.**
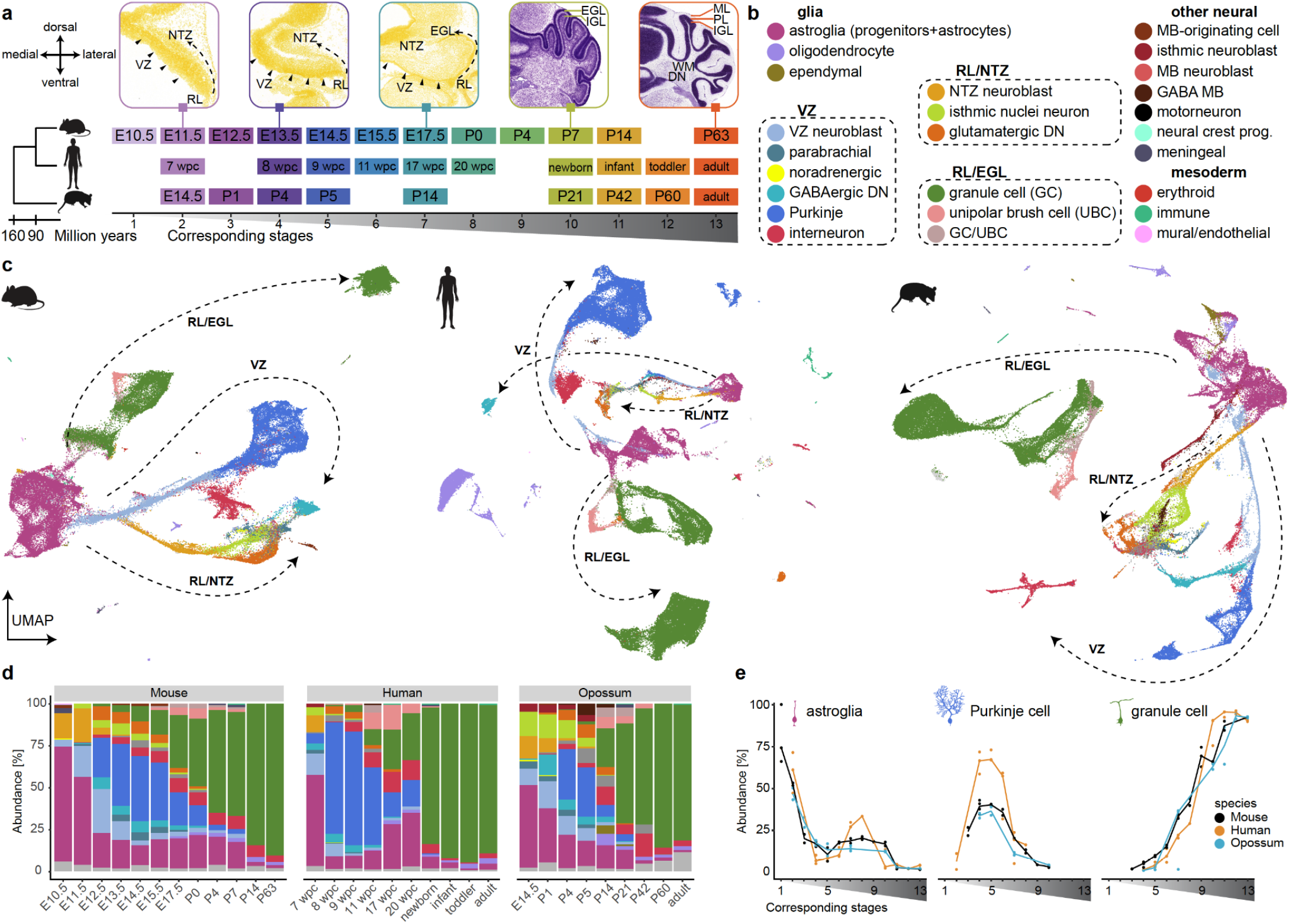
Atlases of cerebellum development across mammals. **a**, Stages sampled in mouse, human and opossum, aligned according to their correspondences across species. Top: coronal sections of the mouse cerebellum^15^ stained with HP Yellow or Nissl. **b**, Cell types detected in this study, grouped based on their developmental origin. Cerebellar neuron types are circled. **c**, Uniform Manifold Approximation and Projection (UMAP) of 115,282 mouse, 180,956 human and 99,498 opossum cells coloured by cell type. The broad neuronal lineages are shown with arrows. **d-e**, Relative cell type abundances across developmental stages in the whole datasets (**d**) or amongst the cerebellar cells (**e**). In **c-d** cell type colours are as in **b**; cells not assigned are in grey. In **e** stages are aligned as in **a** and the line indicates the median of biological replicates. DN, deep nuclei; EGL/IGL, internal/external granule cell layer; MB, midbrain; ML, molecular layer; NTZ, nuclear transitory zone; PL, Purkinje cell layer; RL, rhombic lip; VZ, ventricular zone; WM, white matter.

To establish correspondences between the developmental stages sampled in mouse, human, and opossum, we performed stage-wise cross-species comparisons of (i) synthetic bulk transcriptomes using Spearman correlations of orthologous variable gene expression, (ii) pseudoages^18^ based on the median age of neighbouring mouse cells in the cross-species integrated manifold, and (iii) cellular composition by measuring similarities at the level of cell states (Extended Data Fig. 2d-f). By combining these approaches, we infer, for instance, that the cerebellum of a newborn human corresponds to that of a one week old mouse and a three week old opossum (Fig. 1a). The cerebellar stage correspondences (Fig. 1a) are in agreement with correspondences estimated for somatic organs^19^, which suggests that there is no major heterochrony in the development of the cerebellum across mammals.

Based on the expression patterns of orthologous genes that are differentially expressed within each of the three species, we created a consensus classification of the cellular diversity in the developing mammalian cerebellum (Extended Data Figs. 2c, 3-6). UMAP embeddings of the mouse, human, and opossum datasets show a radiation of lineage-committed cells stemming from a population of proliferating neuroepithelial and radial glial progenitors (arrows in Fig. 1c, Extended Data Fig. 7a). Along the glial and neuronal trajectories, cells are ordered according to their developmental stages (Extended Data Fig. 1e). The major neuronal trajectories (Fig. 1b,c) reflect the known germinal zones in the developing cerebellum: the ventricular zone (VZ), which produces cerebellar GABAergic neurons; the early rhombic lip, which gives rise to glutamatergic neurons assembling at the nuclear transitory zone (RL/NTZ); and the late rhombic lip, which is associated with a secondary germinal zone in the external granule cell layer (RL/EGL)^6^. The VZ cell populations emerge in a defined temporal order, and include parabrachial neurons (marked by *LMX1B* and *LMX1A* expression) and a small group of noradrenergic neurons (*LMX1B, PHOX2B*), both of which migrate to the brainstem during development^20^, as well as all cerebellar GABAergic neuron types; that is, GABAergic deep nuclei neurons (*SOX14*), Purkinje cells (*SKOR2*), and GABAergic interneurons (*PAX2*) (Figure 1b,c, Extended Data Fig. 3a-d). The RL/NTZ cells comprise extra-cerebellar isthmic nuclei neurons that locate to the anterior NTZ during development, and glutamatergic deep nuclei neurons (Extended Data Fig. 4a,b,d,f). The RL/EGL trajectory includes granule cells (GCs) and unipolar brush cells (UBCs) transitioning from progenitors (*ATOH1*) and differentiating cells (*PAX6*) towards defined GC (*GABRA6*) and UBC (*LMX1A*) states (Extended Data Fig. 5a,b,d,e). Along all major neuronal trajectories, cells from different cell types often cluster together at the earliest differentiation states and are designated as VZ neuroblasts, NTZ neuroblasts, and GC/UBC (Fig. 1b,c, Extended Data Fig. 3-5). This clustering is consistent with the observation of a pan-neuronal transcriptional state among the early neuroblasts across the whole developing mouse brain^18^. Further dissection of the VZ neuroblasts, often based on developmental stage (Methods), revealed differential expression of known marker genes of the VZ-derived cell types (e.g., parabrachial and noradrenergic neuron marker *LMX1B*^*20*^ in the early neuroblasts, interneuron marker *PAX2*^*6*^ in the late neuroblasts; Extended Data Fig. 3e,f), consistent with these cells already being lineage-committed, despite common differentiation programs.

The neuroepithelial and radial glial progenitors with temporally progressing transcriptional states, glioblasts, and astrocytes (*SLC1A3, AQP4*) form the most abundant glial lineage in the three datasets (Fig. 1b,c, Extended Data Fig. 6a-f). We refer to the cells in this lineage collectively as astroglia. The oligodendrocyte lineage splits into proliferating oligodendrocyte progenitor cells (OPCs, *PDGFRA*), committed oligodendrocyte precursors (*TNR*), and postmitotic oligodendrocytes (*MAG*; Extended Data Fig. 6a-d,f). Additionally, we detected cells intermediate between astroglial progenitors and OPCs that likely represent the preOPC state (*EGFR*)^18^. preOPCs are relatively abundant in opossum, rare in human, and are not detected in mouse (Extended Data Fig. 6c,d,f), in accordance with the extra-cerebellar origin of most oligodendrocytes in mouse^21^. Ependymal cells (*SPAG17*) are present in small numbers in the mouse and opossum datasets, but were not reliably distinguishable in the human dataset (Extended Data Fig. 6a-d, f). In opossum, we identified a population of ependymal progenitors that is transcriptionally related to glioblasts, but expresses *SPAG17*, which is involved in ciliogenesis (Extended Data Fig. 6a-d,f). We also identified a few neural populations from brain regions adjacent to the cerebellum, resulting from the migration of a group of midbrain-originating cells (*LEF1*) to the cerebellar primordia^12^ or sample contamination (isthmic neuroblasts, midbrain neuroblasts, GABAergic midbrain cells and motor neurons; Fig. 1b,c). We additionally detected neural crest- and mesoderm-derived cell types: meningeal, erythroid, immune (mostly microglia), and vascular (mural and endothelial) cells (Fig. 1b,c).

A comparison of cell type abundances across development revealed highly dynamic temporal patterns that are similar in the three species (Fig. 1d, Extended Data Fig. 7b) and are consistent with the current understanding of cerebellum development^6^. Astroglia (progenitors) are most abundant at the earliest developmental stages (50-56% at stages corresponding to embryonic day E11.5 in mouse), Purkinje cell relative abundances peak at the transition from embryonic to fetal development (E13.5-E15.5 in mouse), and granule cells dominate at late developmental stages, outnumbering all other cell types already in postnatal day P4 mouse, newborn human, and P21 opossum (Fig. 1d,e). We note that from 17 wpc (weeks post conception) onwards, we could only sample parts of the human cerebellum (Supplementary Table 1). The data for these stages might therefore not precisely reflect the cell type proportions in the entire cerebellum, akin to the adult human samples collected from different lobes of the cerebellar cortex or deep nuclei (Extended Data Fig. 7c). By examining the dynamics of individual cerebellar cell types across matched developmental stages, we observed a ∼2-fold higher relative abundance of Purkinje cells in human compared to mouse and opossum at the stages when their relative abundances peak during development (Fig. 1e). The difference remained statistically significant even when additionally considering the VZ neuroblasts (hierarchical Bayesian model; Extended Data Fig. 7d,e). This change in Purkinje cell dynamics in the human lineage could potentially be related to differences in developmental durations between humans and the other mammals and/or the unique presence of basal progenitors in the human cerebellum^9^ that may serve as an additional pool of Purkinje cell progenitors.

Altogether, our snRNA-seq atlases provide a comprehensive view of the cerebellar cell types in mammals and show that their developmental dynamics have been largely conserved for at least 160 million years, except for human Purkinje cells, which have considerably increased their numbers at their abundance peak compared to mouse and opossum.

### Spatiotemporal cell type diversification

Although traditionally viewed as a brain region with a simple cellular architecture, the adult cerebellum is increasingly recognized to exhibit regional specialization of cell types and a complex pattern of functional compartments organized around the parasagittal ALDOC (zebrin II) positive and negative Purkinje cell domains^6–8^. To characterize the molecular diversification of cerebellar cells during development, we examined the within-cell-type heterogeneity in our snRNA-seq atlases. Mouse Purkinje cells could be divided into four developmental subtypes that emerge from VZ neuroblasts with a shared transcriptomic program (Fig. 2a, Extended Data Fig. 3g). Combinatorial expression of transcription factor (TF) *Ebf1* and *Ebf2* genes differentiates the subtypes along the spatial and temporal axes: Purkinje subtypes that locate medially in the developing cerebellum (named by their marker genes as RORB and CDH9 types) display higher *Ebf1* levels than the lateral subtypes (FOXP1, ETV1), whereas *Ebf2* is upregulated in the late-born subtypes (CDH9, ETV1) compared to the early-born subtypes (RORB, FOXP1) (Fig. 2a-c, Extended Data Fig. 3h). The genes variable across the Purkinje developmental subtypes are enriched for the cadherin family of adhesion molecules (homophilic cell adhesion, *P* < 10^−15^), supporting their proposed role in providing a molecular code for the formation of Purkinje cell domains^22^. Based on key marker genes and the correlation of orthologous variable gene expression, we identified the same four developmental Purkinje subtypes in opossum, whereas in human we distinguished two subtypes (*EBF1/2*-low and -high) with only partially defined correspondences to the mouse subtypes (Fig. 2b,d, Extended Data Fig. 3g), possibly related to the differences in human Purkinje cell developmental dynamics and/or sampling. To link the developmental Purkinje subtypes to the recently described adult subtypes in mouse^7^, we used genes that are differentially expressed both among the developmental and adult subtypes to calculate the correlation coefficients between the expression profiles. Despite the gap in Purkinje cell sampling between the two datasets, we detected similarities between early-born subtypes (RORB, FOXP1) and adult *Aldoc*-positive subtypes that are enriched in hemispheric lobules of the cerebellum, late-born medial subtype (CDH9) and *Aldoc*-positive subtypes enriched in posterior vermal lobules; and late-born lateral subtype (ETV1) and *Aldoc*-negative adult subtypes (Extended Data Fig. 3i). Altogether, these results suggest that Purkinje cells with distinct settling patterns are specified not only by their birthdate^6,23^ but also by birthplace.

**Fig. 2.**
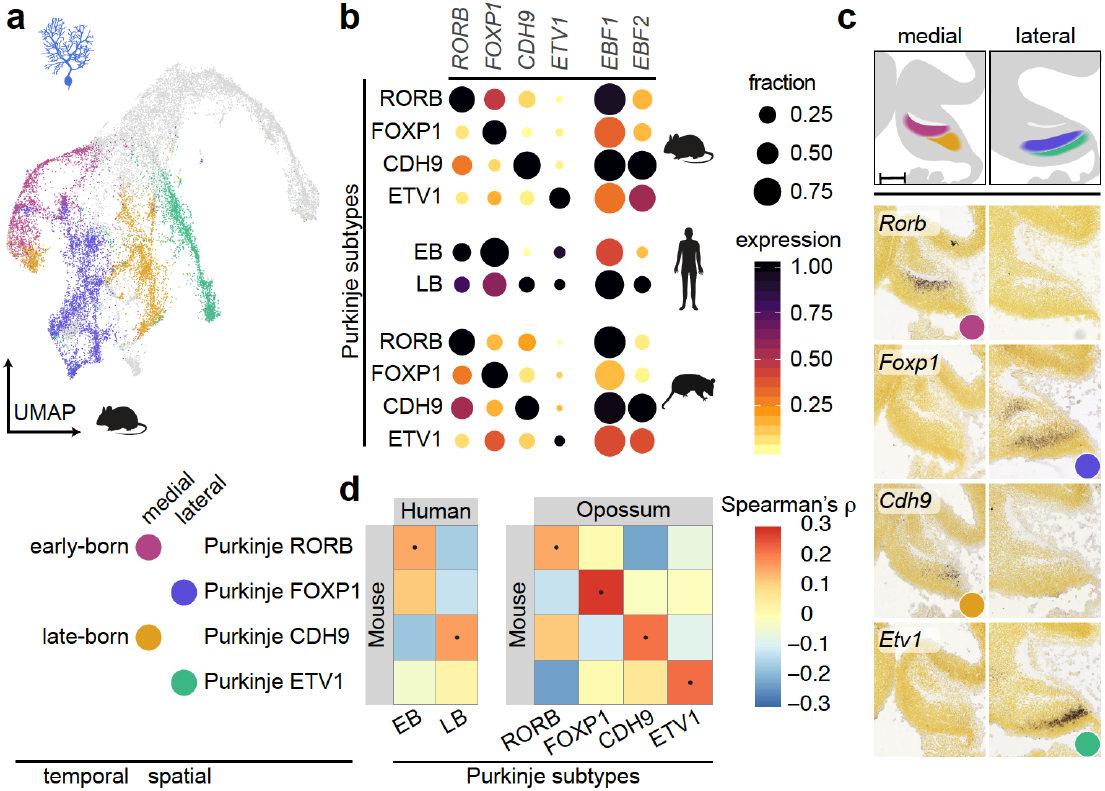
Spatiotemporally defined Purkinje cell subtypes. **a**, UMAP of 23,255 mouse cells assigned to the Purkinje cell lineage. Developmental subtypes are highlighted with colours. **b**, Expression of key marker genes in the Purkinje subtypes in mouse, human and opossum. Dot size and colour indicate the fraction of cells expressing each gene and the mean expression level scaled by species and gene, respectively. **c**, Spatial distribution of Purkinje subtypes in mouse E13.5 cerebellar primordium based on RNA *in situ* hybridisation data^15^ for subtype marker genes. Medial and lateral sagittal sections counterstained with HP Yellow are shown. Scale bar: 200 μm. **d**, Spearman’s correlation coefficients between orthologous variable gene (n=107) expression profiles from mouse, human and opossum Purkinje subtypes. Dots indicate the highest correlation for each column. EB, early-born; LB, late-born.

We further defined 16 subtypes among the other neuronal cell types, 13 of which were detected in all three species (Extended Data Figs. 3-5). GABAergic interneurons comprise a set of five homologous interneuron subtypes and an unknown cell group (*MEIS2)* in opossum (Extended Data Fig. 3j-l,n). The interneuron subtypes shared across the three species follow the known temporal sequence of production^6^ and are directly matched to the transcriptionally-defined subtypes with layer-specific localizations in the adult mouse cerebellum^7^ (Extended Data Fig. 3m). Among RL/NTZ cells in the three species, glutamatergic deep nuclei neurons diversify into two populations that display different spatial and temporal patterns, consistent with previous studies^12,24^, and isthmic nuclei neurons include subsets that express marker genes related to dopaminergic (*NR4A2*), cholinergic (*SLC5A7;* subtype not detected in human), or somatostatin (*SST*) identities (Extended Data Fig. 4c-e,g,h). Consistent with prior work on the adult mouse cerebellum^7^, developing UBCs and GCs display continuous variation in all three species (Extended Data Fig. 5c,f). We classified UBCs into two subsets: one expresses the canonical pan-UBC marker *EOMES* and is co-labeled by markers of known UBC subtypes^6,7^ (*TRPC3, GRM1, CALB2*), whereas the other is a so far uncharacterized *EOMES*-low UBC-like subset that is positive for *HCRTR2*, which marks scattered cells in the internal granule cell layer in the P4 mouse (Extended Data Fig. 5g,h). Differentiating GCs clustered into early and late populations, and in mouse and opossum we additionally detected a distinct *KCNIP4* and *OTX2* expressing subset (Extended Data Fig. 5c,f,g,i), which might not have been distinguished in human due to sampling differences. Comparing these three groups to the GC subtypes defined in the adult mouse cerebellum^7^, we observed correspondences with the adult subtypes that are spatially invariant (early), enriched in the posterior hemisphere (late) or nodulus (KCNIP4) (Extended Data Fig. 5j), supporting the notion that the topographic heterogeneity of granule cells is at least partially driven by the temporal ordering of GC differentiation^25^.

The diversity of neuronal populations in the developing cerebellum is paralleled by the heterogeneity among the progenitor populations (Extended Data Fig. 6a). In the three species, the embryonic neurogenic progenitors display a rostrocaudal gradient of molecular variation along the neuroepithelium and include a group of potentially apoptotic cells (*NCKAP5* low, *BCL2L11* high ; Extended Data Fig. S6b-e). During the next period (corresponding to fetal and early postnatal stages in mouse), the neurogenic progenitors are gradually outnumbered by bipotent progenitors (i.e., producing both interneurons and parenchymal astrocytes^6,26^) and gliogenic progenitors (producing parenchymal and Bergmann astrocytes^27^) that **–** compared to the early progenitors **–** have decreased expression of cell cycle-related genes (Extended Data Fig. 6c,d,f,g). In the three species, we detected two glioblast populations, prospective white matter glioblasts and astroblasts (Fig. S6c,d,f), corroborating our previous observations in the developing mouse cerebellum based on chromatin accessibility data^24^.

Collectively, these results suggest developmental specification of the regional heterogeneity amongst the cerebellar cell types and highlight the overall conservation of the cellular architecture, including neural subtypes, of the developing cerebellum across mammals.

### Cell type-defining transcriptional programs

Having established cross-species correspondences between developmental stages, as well as cell types and states in the cerebellum datasets from the three mammals, we next sought to characterize global gene expression patterns in the three species. We aggregated expression values from all cells into cell type pseudobulks for each sample and performed principal components analysis (PCA) using orthologous genes expressed in all species. The two first principal components order samples by developmental stage and split glial and neuronal cells, while the third component further separates the neuronal types (Fig. 3a, Extended Data Fig. 8a). In a separate PCA only of neurons, the first component orders samples by developmental stage and the second separates neuronal types (Extended Data Fig. 8b,c). These clustering patterns indicate that gene expression variance in the developing cerebella is to a large extent explained by developmental and cell type signals that are shared across the species. Thus, we sought to identify the core gene expression programs that underlie the identity of cerebellar cell types, akin to previous comparative cross-species approaches^28–30^. For this, we called enriched genes for each cell state in each species using the term frequency–inverse document frequency (TF-IDF) transformation and determined their overlap across species. On average, 61% of the enriched genes (markers) in each species are cell state-specific, 26% are enriched in two cell states and 13% in three or more states (Extended Data Fig. 8d). Similarly to observations for the adult motor cortex^28^, many of the markers displayed cell state enrichment in only one species (Fig. 3b,c, Extended Data Fig. 8e). Nevertheless, each cell state category exhibited a set of conserved markers (Supplementary Table 5) that likely represent genes that drive cerebellar cell type identities, given that their cellular expression specificity has been retained for at least 160 million years of evolution. Consistently, conserved markers are associated with pertinent gene ontology terms, including ‘myelination’ for oligodendrocytes and ‘Purkinje cell differentiation’ for Purkinje cells (Supplementary Table 6). The numbers of conserved markers vary between cell states and are generally highest for the most mature states (Extended Data Fig. 8e). The cell states that share conserved markers are usually closely related, either in terms of cell type lineage or maturation status (Extended Data Fig. 8f). At states of differentiation, when cell (sub)type specification is ongoing, there is an enrichment of TF genes among conserved markers (Extended Data Fig. 8e), in line with the central role of TFs in the induction of the downstream effector genes characteristic of a cell type^31^. The list of conserved marker genes across all cell states includes 185 TF genes (Supplementary Table 5, Extended Data Fig. 8g), many of which are known to have functions in specific cerebellar cell types^6,10,32,33^ (e.g., *PAX3* in progenitors, *HOPX* in astrocytes, *ESRRB* and *FOXP2* in Purkinje cells, *PAX2* in interneurons, and *PAX6* and *ETV1* in granule cells). However, this list also includes potential novel regulators, for instance interneuron-enriched *PRDM8* and *BHLHE22*, which have previously been shown to form a repressor complex involved in pallial circuit formation^34^, and *SATB2*, which is transiently upregulated in differentiating granule cells and mainly known as a determinant of neocortical upper-layer neurons^35^ (Extended Data Fig. 8g). Thus, the identified conserved TF code of mammalian cerebellum development provides a shortlist of candidates for the functional elucidation of the mechanisms underlying the specification of the respective cell types.

**Fig. 3.**
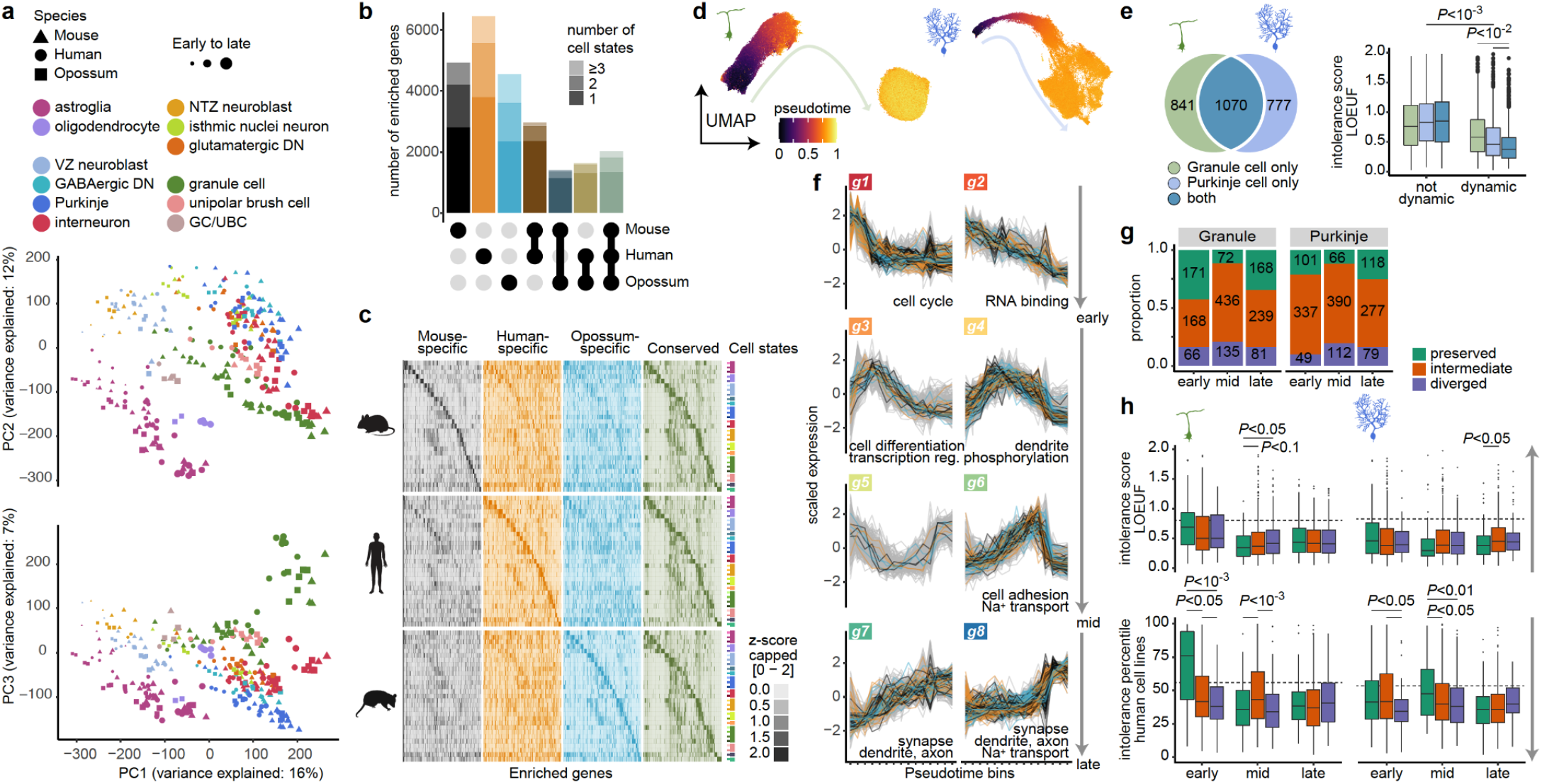
Cell type-defining transcriptional programs. **a**, Principal components analysis based on 10,276 orthologous genes expressed in all species. Data points represent cell type pseudobulks for each biological replicate. **b**,**c**, Numbers (**b**) and expression patterns (**c**) of species-specific and conserved markers. 20 genes per cell state are shown in **b. d**, UMAPs of cells from granule cell (left) or Purkinje cell (right) lineages aligned across species and coloured by diffusion pseudotime values. **e**, Intolerance to functional mutations in human population (LOEUF, loss-of-function observed/expected upper bound fraction; low values indicate intolerance) for genes dynamic or non-dynamic across differentiation of granule cells, Purkinje cells, or both neuron types in all species. Numbers per category are indicated on the left. **f**, Clusters (g1-g8) of gene expression trajectories during granule cell differentiation, ordered from early to late differentiation based on the mean center-of-mass values of the confident cluster members’ trajectories. Strongly preserved trajectories of the orthologues are highlighted. Examples of enriched gene ontology categories for the genes with preserved trajectories are indicated. **g**, Proportion of human genes with strongly preserved, intermediate and diverged trajectories in early, mid (excluding g5) and late clusters. **h**, Intolerance to functional mutations in human population (top; LOEUF) or cell lines (bottom) for human genes from different trajectory conservation groups across early, mid and late clusters of granule cell (left) or Purkinje cell (right) differentiation. The arrows point towards more tolerance and dashed lines indicate the median score of the non-dynamic genes. In **e** and **h**, adjusted *P*-values were calculated via a permutation test of pairwise comparisons between all categories (**e**) or trajectory conservation groups (**h**); boxes represent the interquartile range, and whiskers extend to extreme values within 1.5 times the interquartile range from the box.

The above analyses are based on discrete categories of cell states, but developmental processes are inherently continuous. Thus, we set out to also delineate the conserved gene expression cascades across differentiation of the principal cerebellar neuron types **–** Purkinje and granule cells. We integrated cells belonging to each of the two neuronal lineages from all species and performed diffusion pseudotime^36^ analyses (Fig. 3d, Extended Data Fig. 9a). Corresponding cell states across species display comparable pseudotime values (Extended Data Fig. 9b), and the distribution of the values across stages (Extended Data Fig. 9c) is in accordance with the different production modes of the two neuron types **–** transient for Purkinje cells and protracted for granule cells^6,37^ **–** thus corroborating the alignment of cells across species and stages. Next, for both neuron types, we identified orthologous genes with dynamic gene expression throughout differentiation in all three species (Methods). For each neuron type, 56-58% of the dynamic genes were shared with the other type, suggesting considerable overlap in their differentiation programs (Fig. 3e). We then characterized the set of dynamic genes associated with the two neuron types using different metrics of functional constraints: the *in vivo* gene essentiality scores that measure tolerance to heterozygous inactivation in human population, and the *in vitro* score that is inferred from viability of human cell lines^38,39^. We found that dynamic genes are enriched for *in vivo* essential genes, but not for *in vitro* essential genes, likely because the latter include many genes involved in the cell cycle and cellular metabolism^40^ (Fig. 3e, Extended Data Fig. 9d). In line with studies linking phenotypic severity to expression pleiotropy^19,41^, we found that the dynamic genes shared between the two neuron types are under stronger constraint than genes dynamic in one cell type only (Fig. 3e, Extended Data Fig. 9d). Dynamic genes are additionally enriched for TFs and genes associated with inherited developmental diseases affecting the nervous system^42^ (Extended Data Fig. 9e,f). We further focused on neurodevelopmental and neurodegenerative diseases^13^ and malignancies^43^ that are directly linked to cerebellar functions and cell types. Genes associated with cerebellar malformations (Dandy-Walker malformation and hypoplasia), spinocerebellar ataxia, and medulloblastoma are enriched among the set of dynamic genes shared between the two neuron types, whereas high-confidence risk genes of autism spectrum disorder and intellectual disability are additionally enriched among conserved genes dynamic during Purkinje cell differentiation only (Extended Data Fig. 9f). These results indicate that many of the cerebellar disease-linked genes likely affect more than one neuron type.

Next, we grouped the genes dynamic across neuronal differentiation in clusters based on their expression trajectories, and determined center-of-mass values for the individual trajectories within each cluster to ensure comparable distributions across species (Fig. 3f, Extended Data Fig. 9g,h). By comparing the cluster assignments of the orthologues from each species, we assigned the genes into three trajectory conservation groups: (i) 23% of genes, on average, were defined as strongly preserved with orthologues confidently assigned (*cluster membership* > 0.5 and *P* > 0.5) to the same trajectory cluster; (ii) 17% of genes were defined as diverged, based on the differential cluster assignment (*P* < 0.05) of at least one of the orthologues; (iii) the remaining 60% of genes were defined as having intermediate trajectory conservation (Fig. 3g). Consistently, the maximum distance between the trajectories of the orthologues progressively decreases from the most strongly to the least preserved trajectory group (Extended Data Fig. 9i). Among the strongly preserved trajectory genes of granule and Purkinje cells, there are 30 and 43 TF genes, respectively, including several TFs with well-characterized roles in neuronal differentiation in the cerebellum (e.g., *PTF1A, RORA* for Purkinje cells; *PAX6, ETV1* for granule cells^6^). We ranked the TFs based on the center-of-mass values of their clusters and individual trajectories for both neuron types, and confirmed the expression patterns of many of the TFs using mouse RNA *in situ* hybridisation data^15^ (Extended Data Fig. 9j,k). Thus, these analyses reveal a conserved program of TFs, the expression of which follows closely matched patterns during Purkinje or granule cell differentiation in the three mammals. Genes with strongly preserved trajectories are under higher levels of functional constraints in humans, but the mode of essentiality depends on the expression patterns. Genes expressed early during neuronal differentiation show enrichment of genes essential for cellular viability (*in vitro*), consistent with their involvement in cell cycle and/or RNA metabolism, whereas the mid and/or late-expressed genes show low tolerance to heterozygous inactivation in humans (*in vivo*), and have functions in differentiation and neuronal maturation (Fig. 3f,h, Extended Data Fig. 9g,l). Altogether, the pivotal role of the genes with strongly preserved trajectories during granule and Purkinje cell differentiation between mammals is substantiated by the associated viability phenotypes.

### Evolutionary change in gene expression

Species-specific phenotypic features emerge during evolution through changes in gene expression programs. We therefore aimed to systematically identify genes that display distinct expression patterns in cerebellar cells in one of the three species. First, we traced genes with diverged expression trajectories across species along the Purkinje or granule cell differentiation paths (Fig. 3g). Using opossum as an evolutionary outgroup, we assigned the trajectory changes to the mouse or human lineage (i.e., polarized the changes; Fig. 4a). In granule cells, we found a relative excess of trajectory changes in the human lineage (136 in human, 54 in mouse; *P* < 10^−6^, binomial test), whereas in Purkinje cells, we found similar numbers of changes in the human (47) and mouse (42) lineages (Fig. 4b, Supplementary Table 7). In each lineage, only a few (1-4) genes have changed trajectories in both cell types, suggesting that changes in regulatory programs are largely cell type-specific. The trajectory changes include shifts in both directions along the differentiation path (towards less or more mature states) and involve all types of differentiation trajectories (Fig. 4a, Extended Data Fig. 10a,b). We attempted to obtain a quantitative measure of the amount of change for each gene by assessing the maximum and minimum pairwise distances between the trajectories of orthologues from the three species (Methods). This approach identified *SNCAIP* (synuclein alpha interacting protein) and *MAML2* (a transcriptional coactivator in NOTCH-signaling pathway) as having evolved the strongest changes in expression trajectories during granule cell and Purkinje cell differentiation, respectively, in the human lineage (Fig. 4c,d, Extended Data Fig. 10c,d).

**Fig. 4.**
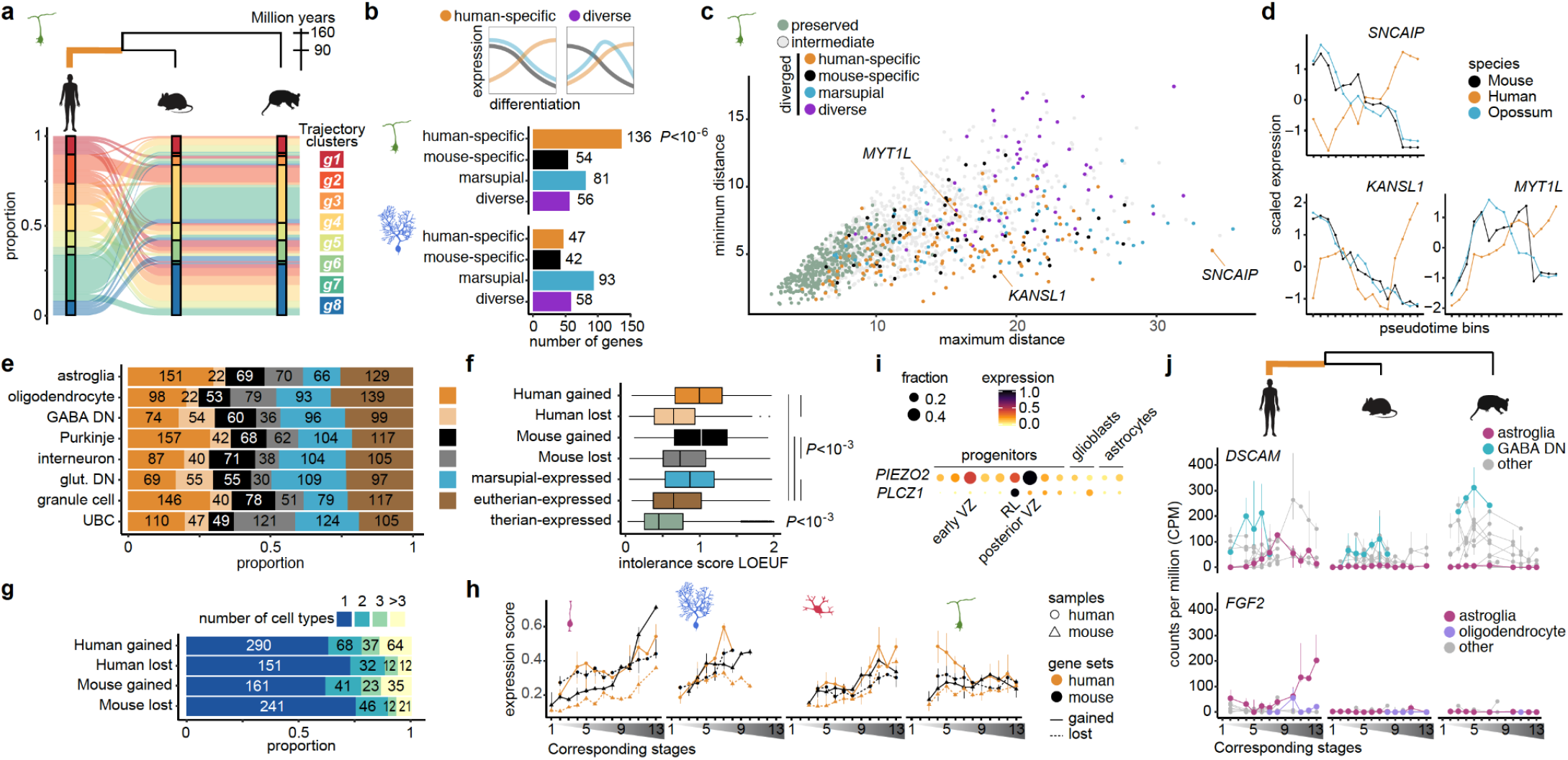
Evolutionary change in gene expression. **a**, Changes in gene expression trajectories during granule cell differentiation assigned to the human lineage. For each species, trajectory clusters are ordered from early to late differentiation. **b**, Numbers of genes with trajectory changes during granule cell (middle) or Purkinje cell (bottom) differentiation in different phylogenetic branches. The scheme (top) shows examples of a change in the human lineage and a diverse pattern that cannot be assigned to a lineage. **c**, Scatter plot of minimum and maximum pairwise distances between the trajectories of orthologues from the three species. High maximum and low minimum distances indicate the strongest lineage-specific trajectory changes. **d**, Examples of genes that evolved a new trajectory during granule cell differentiation in the human lineage. **e**, Distribution of presence/absence expression differences assigned to different phylogenetic branches in the cerebellar cell types. **f**, Intolerance to functional mutations in human population (LOEUF; low values indicate intolerance) for genes grouped based on the presence/absence of expression in the three therian mammals. The summarised values from all eight cell types included in this analysis are shown; boxes represent the interquartile range, and whiskers extend to extreme values within 1.5 times the interquartile range from the box. **g**, Number of cell types in which genes gained or lost expression in different phylogenetic branches. **h**, Expression of genes with gained or lost expression in the mouse or human lineage in astroglia, Purkinje cells, interneurons, and granule cells across development. The expression of the genes that were lost in the human lineage were evaluated in the mouse samples, and *vice versa*. **i, j**, Examples of genes that gained expression in astroglial cells in the human lineage. *PIEZO2* and *PLCZ1* are enriched in human posterior ventricular zone or rhombic lip progenitors. *DSCAM* is expressed in GABAergic deep nuclei neurons in all studied species; *FGF2* is not expressed in any neural cell type in mouse and opossum. In **j**, dots represent the median, bars indicate the range between the biological replicates. In **h** and **j**, stages are aligned across species as shown in Fig. 1a; the line indicates the median of biological replicates. In **b** and **f**, adjusted *P*-values were calculated via a binomial (**b**) or permutation test of pairwise comparisons (**f**). In **i**, dot size and colour indicate the fraction of cells expressing each gene and the scaled mean expression level. GABA DN, GABAergic deep nuclei neurons; glut DN, glutamatergic deep nuclei neurons; RL, rhombic lip; UBC, unipolar brush cell, VZ, ventricular zone.

We next sought to identify genes with an even more fundamental expression change; that is, genes that display presence/absence expression differences between species in one or several of the cerebellar cell types (Methods). Out of the 9,189 orthologous genes included in this analysis, 3,586 displayed presence/absence expression differences in at least one cell type. These differences are consistent with the expression levels of the affected genes in mouse, human, and opossum cerebellum development, as inferred from bulk RNA-sequencing data^19^ (Extended Data Fig. 10e). After polarizing the differences between mouse and human, we found, on average, 112 gains and 40 losses in the human lineage, and 63 gains and 61 losses in the mouse lineage per cell type (Fig. 4e, Supplementary Table 7). Compared to the genes expressed in all species, genes that gained expression in the human or mouse lineage are under weaker functional constraint, whereas the genes that lost expression show intermediate levels of constraint (Fig. 4f, Extended Data Fig. 10f). Most expression gains and losses are cell-type specific (Fig. 4g, Extended Data Fig. 10g,h). Nevertheless, cell type-specific expression gains are more likely to involve genes that were already expressed in other neural cell types in the cerebellum, than genes newly recruited to the cerebellar transcriptomes (Extended Data Fig. 10i). Among the newly incorporated genes in the human cerebellum are *ZP2*, a zona pellucida glycoprotein gene, and *CPLX4*, a complexin that is expressed in the mammalian retina and functions in synaptic vesicle exocytosis^44^. We detected *ZP2* expression in mature granule cells and glutamatergic deep nuclei neurons, and *CPLX4* in granule cell layer interneurons, patterns that are supported by human immunohistochemistry data^45^ (Extended Data Fig. 10j). Based on adult bulk RNA-sequencing data from nine mammals (including six primates)^46^, we inferred that *ZP2* expression in the adult cerebellum was acquired specifically in human in the past ∼7 million years, after the human-chimpanzee split, in line with previous findings^47^, and that the distinct *CPLX4* expression emerged in the lineage leading to the great apes (Extended Data Fig. 10j). Both of these genes are expressed late in development. To understand if expression in late development is a general pattern for genes that have gained or lost expression in the mouse or human lineage, we assessed their expression patterns in the respective cell type during development, and across cell states and subtypes (genes not expressed in the human lineage were evaluated in the mouse, and *vice versa*). This analysis revealed that the aggregated expression levels of these genes increase during development in most cell types (Fig. 4h, Extended Data Fig. 10k). A notable exception occurs in human granule cells, which express the genes that gained expression in the human lineage at high levels already at early developmental stages, and at the granule cell progenitor state (Fig. 4h, Extended Data Fig. 10l). Compared to mouse progenitor cells (astroglia), we observed relatively high levels of gene expression gains in the human rhombic lip and posterior ventricular zone progenitor subtypes that are present at 8-11 wpc (Extended Data Figs. 10l, 6c). Among the genes with gained expression in human astroglia, 19 genes are enriched in the latter populations of human progenitors (TF-IDF, hypergeometric test, *P* < 0.01). These genes are enriched for roles in the extracellular compartment and signal transduction (binomial test, *P* < 0.05), and include the mechanosensitive ion channel *PIEZO2* and the phospholipase *PLCZ1* expressed in human posterior ventricular zone or rhombic lip progenitors, respectively (Fig. 4i, Supplementary Table 8). Thus, we suggest that this gain of expression could play a role in the specification of the unique pool of basal progenitors identified in the developing human cerebellum^9^.

We then asked whether genes with trajectory or presence/absence expression differences between human and mouse include genes associated with cerebellum-linked diseases. This analysis revealed 49 disease genes (Supplementary Table 7). Among the disease-associated genes with trajectory differences is the aforementioned *SNCAIP* (Fig. 4d), which is frequently duplicated in group 4 medulloblastoma^48^, a childhood brain tumour that, notably, has been difficult to model in mouse^49^. Twelve genes associated with autism spectrum disorder and/or intellectual disability have different expression trajectories during granule cell (e.g., *MYT1L* and *KANSL1*; Fig. 4d) or Purkinje cell differentiation (e.g. *SMARCA2, DIP2C* and *FOXP1*; Extended Data Fig. 10d). 25 disease-associated genes have gained or lost expression in at least one of the cerebellar cell types in mouse or human; for instance autism and Down syndrome-associated *DSCAM* gained expression in human astroglia, and *FGF2*, implicated in pilocytic astrocytoma, gained expression in human astroglia and oligodendrocytes (Fig. 4j).

Altogether, our comparative molecular analyses of mammalian cerebellar development unveiled numerous candidate genes with potentially adaptive expression changes, and disease genes for which functional characterization in a mouse model might not reflect all disease manifestations in human. Thus, these results underline the importance of studying human development to understand human disease, an approach we have taken in our companion paper to elucidate the origins of childhood brain tumours (Okonechnikov *et al*.).

## DISCUSSION

In this study, we used a comprehensive comparative approach to characterize the evolution of the mammalian cerebellum and its development from the beginning of neurogenesis to adulthood. Our snRNA-seq atlases of ∼400,000 cells from the mouse, human, and opossum cerebellum (https://apps.kaessmannlab.org/sc-cerebellum-transcriptome) revealed largely shared gene expression programs across homologous cell types. Based on these conserved patterns, we provide a catalogue of genes associated with the specification and maintenance of cerebellar cell type identities across at least ∼160 million years of mammalian evolution.

Identifying the genes that establish and maintain the diversity of cerebellar cell types is particularly important because of the growing evidence for a pronounced molecular heterogeneity within the seemingly uniform cellular organization of the mature cerebellum^7,25,37^. Our analyses of the developing cerebellum support early spatiotemporal patterning of Purkinje and granule cell subtypes that contributes to the cellular organization in the adult. We found consensus subtypes for most neural cell types in the developing cerebellum and observed equivalent sequences and timings of cell type emergence in the three species, in line with the previously proposed conservation of the cerebellum’s cellular architecture and developmental program across amniotes^4,50^. A notable exception is the significantly higher relative abundance of Purkinje cells during early fetal development in human. This occurs at the stages when Purkinje cells are generated and is consistent with the expansion of neuronal progenitor pools in the human cerebellum in the same time window^9^. Given that Purkinje cell signals regulate the transit amplification of granule cell progenitors^6,37^, we suggest that higher numbers of Purkinje cells could augment the generation of granule cells and lead to the increase in cerebellar cell numbers required to match the expansion of the neocortex in the human lineage^3^.

Evolutionary innovation in cellular programs is expected to be driven by lineage-or species-specific differences in gene expression. Consistently, we identified a set of genes that are recruited to the transcriptomes of subpopulations of human progenitor cells in the cerebellar germinal zones, potentially underlying their human-specific characteristics, which allow these cells to remain proliferating outside of the ventricular zone^9^. Furthermore, we found presence/absence expression differences between the three mammals for all neural cell types, and detected shifts in the gene expression trajectories during Purkinje and granule cell differentiation. Both of these types of expression changes are highly cell type-specific, indicating that concerted shifts in gene expression across many cell types are rare. This cell type-specific gene expression divergence during evolution is likely facilitated by the modular organization of gene regulatory control^51^ and the high cell-type specificity of *cis*-regulatory elements^24^. In most cerebellar cell types, the genes that gained or lost expression in the human and mouse lineages are more active at later developmental stages, consistent with the progressively increasing molecular divergence of the cerebellum (and other organs) between species during development due to overall decreasing purifying selection that facilitates adaptations driven by positive selection^19,52^. The potential functional relevance of the individual expression shifts to interspecies phenotypic differences requires further interrogation. Notably, several heterochronic gene expression adaptations were previously associated with profound phenotypic effects: SATB2 was shown to contribute to the differences in cortical wiring between eutherians and marsupials^53^, NEUROD1 to the presence of granule cells’ transit amplification in amniotes and absence in amphibians^54^, and LHX9 to the development of different numbers of cerebellar deep nuclei in birds and mammals^55^.

Altogether, our study provides a comprehensive map of the cellular and molecular diversity of the cerebellum across development and evolution for three mammalian species. We used this comparative map to create a resource of candidate gene sets that underlie core ancestral gene expression programs of cell fate specification in the cerebellum as well as genes that contribute to phenotypic innovations of this brain structure during evolution. In our companion paper (Okonechnikov *et al*.), we used the human cerebellum atlas to elucidate the cellular origins of childhood brain tumours, highlighting how our multidimensional dataset can be further leveraged to advance a mechanistic understanding of brain evolution, development, and disease.

## METHODS

### Data reporting

No statistical methods were used to predetermine sample size. The experiments were not randomized and investigators were not blinded to allocation during experiments and outcome assessment.

### Sample collection and ethics statement

The human prenatal samples used in this study were donated voluntarily to the MRC Wellcome Trust Human Developmental Biology Resource (HDBR; UK) by women who had an elective abortion and had given written informed consent to donate fetal tissues for research. The prenatal samples had normal karyotypes and were classified as belonging to a particular Carnegie stage or week post conception (wpc) according to their external physical appearance and measurements. The human postnatal samples are from the University of Maryland Brain and Tissue Bank of National Institutes of Health NeuroBioBank (USA), Chinese Brain Bank Center (CBBC) in Wuhan and Lenhossék Human Brain Program, Human Brain Tissue Bank at Semmelweis University (Hungary). Informed consent for the use of tissues for research was obtained in writing from donors or their family. All postnatal samples come from healthy non-affected individuals defined as normal controls by the corresponding brain bank. The use of human samples was approved by an ERC Ethics Screening panel (associated with H.K.’s ERC Consolidator Grant 615253, OntoTransEvol) and ethics committees in Heidelberg (authorization S-220/2017), North East-Newcastle & North Tyneside (REC reference 18/NE/0290), London-Fulham (REC reference 18/LO/0822), Ministry of Health of Hungary (No.6008/8/2002/ETT) and Semmelweis University (No.32/1992/TUKEB).

RjOrl:SWISS time-mated pregnant mice (*Mus musculus*), litters at postnatal days 0-14 and adult mice were purchased from Janvier Labs (France). Gray short-tailed opossums (*Monodelphis domestica*) were bred in a colony in Museum für Naturkunde Berlin, Leibniz Institute for Research on Evolution and Biodiversity Science at the Humboldt University Berlin (Germany) or Texas Biomedical Research Institute (USA; Supplementary Table 1). Mouse and opossum stages were dated according to the time of copulation (E, embryonic day) or birth (P, postnatal day). The adult mice were sacrificed by cervical dislocation, the pups by decapitation. Opossums were sacrificed by isoflurane overdose. All animal procedures were performed in compliance with national and international ethical guidelines for the care and use of laboratory animals, and were approved by the local animal welfare authorities: Heidelberg University Interfaculty Biomedical Research Facility (T-63/16, T-64/17, T-37/18, T-23/19), Vaud Cantonal Veterinary Office (No.2734.0) and Berlin State Office of Health and Social Affairs, LAGeSo (T0198/13).

### Dissections

Mouse and opossum cerebella were dissected as whole or in 2 halves as described^24^. Mouse E10.5 and E11.5 and opossum E14 cerebella were pooled from 2-3 littermates. (Supplementary Table 1). For human 7-11 wpc samples whole cerebellar primordia were collected and divided into halves or representative fragments (ca 25% of the primordium cut perpendicular to its long axis; Supplementary Table 1). For human 17-20 wpc samples fragments of cerebellum were collected. Newborn, infant and toddler samples include fragments of cerebellar cortex. Adult human samples were dissected from the posterior lobe of the cerebellar hemispheres (crus I and II), vermis (lobules VI-VIII), flocculonodular lobe and dentate nucleus using the micropunch technique^56^. Developmental samples were kept in ice-cold PBS during the dissection, most of the meninges were removed and the samples were snap-frozen in liquid nitrogen and stored at -80 °C. If available, samples from both sexes were used for data production.

### Preparation of nuclei

The nuclei were extracted as described^57^ with modifications. Briefly, the frozen tissue was homogenized on ice in 250 mM sucrose, 25 mM KCl, 5 mM MgCl_2_, 10 mM Tris-HCl (pH 8), 0.1% IGEPAL, 1 µM DTT, 0.4 U/µl Murine RNase Inhibitor (New England BioLabs), 0.2 U/µl SUPERase-In (Thermo Fisher Scientific), cOmplete Protease Inhibitor Cocktail (Roche) by trituration and/or using a micropestle. After 5 minutes of incubation the remaining bits of unlysed tissue were pelleted by centrifugation at 100g for 1 minute at 4°C. The cleared homogenate was centrifuged at 400g for 4 minutes to pellet the nuclei. Supernatant at this step was collected as cytoplasm extract. Nuclei were washed 1-2 times in the homogenization buffer, collected by centrifugation and resuspended in 430 mM sucrose, 70 mM KCl, 2 mM MgCl_2_, 10 mM Tris-HCl (pH 8), 0.4 U/µl Murine RNase Inhibitor, 0.2 U/µl SUPERase-In, cOmplete Protease Inhibitor Cocktail (that optionally allowed storage of the prepared nuclei at -80°C). If needed, the nuclei were strained using 40 µm Flowmi strainers (Sigma). For estimation of nuclei concentration, Hoechst DNA dye or propidium iodide was added and nuclei were counted on Countess II FL Automated Cell Counter (Thermo Fisher Scientific). Hoechst-positive nuclei from adult human vermis, flocculonodular lobe and deep nuclei were FACS-sorted in PBS to remove cellular debris present in preparations from white matter-rich brain tissues. A few modifications in the nuclei preparation method were applied in the pilot phase of data production: (1) the tissue was lysed in HEPES-based homogenisation buffer (250 mM sucrose, 25 mM KCl, 5 mM MgCl_2_, 10 mM HEPES (pH 8), 0.1% IGEPAL, 1 µM DTT, 0.4 U/µl Murine RNase Inhibitor, 0.2 U/µl SUPERase-In), the nuclei were fixed with DSP (dithio-bis[succinimidyl propionate]) as described^58^ and then processed as above; (2) the tissue was lysed in Triton X-100 homogenisation buffer (320 mM sucrose, 5 mM CaCl_2_, 3 mM Mg-acetate, 10 mM Tris-HCl (pH 8), 0.1 % Triton X-100, 2 mM EDTA, 0.5 mM EGTA, 1 mM DTT, 0.4 U/µl Murine RNase Inhibitor, 0.2 U/µl SUPERase-In). The two mouse datasets produced with the modified protocols were comparable to the ones produced with the standard protocol in terms of cellular composition and gene expression (Extended Data Fig. 1c) and were therefore included in the merged datasets. Supplementary Table 1 lists nuclei preparation details for each sample.

### RNA extraction and sample quality control

Cytoplasm extracts or nuclei suspensions were mixed with 40 mM DTT-supplemented RLT buffer (Qiagen) and 100% ethanol at 2:7:5 ratio. The mixtures were subjected to RNA purification with RNeasy Micro Kit (Qiagen). RNA quality numbers (RQN), determined on Fragment Analyzer (Advanced Analytical), were above 7 for all samples except some of the human infant and toddler samples, and opossum P42 and adult samples (Supplementary Table 1).

### Library preparation and sequencing

Chromium Single Cell 3’ Reagent kits (v2 or v3 chemistry) and the Chromium Controller instrument (10x Genomics) were used for single cell barcoding and library construction according to manufacturer’s protocols. In most of the experiments 15,000 nuclei were loaded per channel (range 13,000-17,000). cDNA was amplified in 12 PCR cycles. Libraries were quantified on Qubit Fluorometer (Thermo Fisher Scientific) and the average fragment size was determined on Fragment Analyzer. Libraries were sequenced on Illumina NextSeq 550 (26/28 cycles for Read 1, 8 cycles for i7 index, 57/56 cycles for Read 2 in case of v2/v3 libraries) or HiSeq 4000 (26/28 cycles for Read 1, 8 cycles for i7 index, 56-74/74 cycles for Read 2 in case of v2/v3 libraries).

### Genome annotations

For human and mouse we used gene annotations from Ensembl release 91^59^, based on reference genome assemblies hg38 (human) and mm10 (mouse). For opossum, we used monDom5-based annotations from Ensembl release 87 extended using stranded poly(A) RNA-seq datasets^19^ from the adult brain as described^60^. We split each of the opossum chromosomes 1 and 2 at the nucleotide position 536,141,000, which does not overlap any gene, given that the large size of these chromosomes is not compatible with many bioinformatics pipelines. We generated custom genome reference files with *cellranger mkref* pipeline from Cell Ranger (10x Genomics). Since snRNA-seq captures both mature mRNA and unspliced pre-mRNA, we additionally created pre-mRNA references by merging exons and introns of each gene. Orthology relationships were extracted from Ensembl release 91. The list of mouse transcription factors was downloaded from the animal TFDB (v.3.0)^61^.

### General data processing and quality control

Raw sequencing data were demultiplexed using *cellranger mkfastq* pipeline (Cell Ranger versions 2 or 3.0.2, 10x Genomics, internally calling *bcl2fastq*). Read alignment and counting of Unique Molecular Identifiers (UMIs) was performed with *cellranger count*. Reads from each library were aligned and counted in two modes: (1) using the mature mRNA reference (exons) or (2) the pre-mRNA reference (exons and introns). We used the pre-mRNA counts (exons and introns) in most of the analyses to maximise the amount of reads for quantification. We used the mature mRNA counts (exons) in specific cases: library-level quality control (Extended Data Fig. 1c) and presence/absence expression differences (Fig. 4e-h,j, Extended Data Fig. 10e-i,k,l).

Valid barcodes (i.e. droplets containing a nucleus) were identified by leveraging the higher abundance of intronic reads (originating from pre-mRNAs) in nucleus-containing droplets compared to empty droplets that mostly contain background levels of cytoplasmic mature mRNAs. For each barcode the fraction of intronic UMI counts was estimated as [1-(exonic counts/pre-mRNA counts)]. Barcodes were clustered by their fraction of intronic UMIs using a Gaussian mixed model (mclust^62^ package v5.4.3, R) with two expected clusters. The barcodes in the cluster with higher mean fraction of intronic reads were considered as valid. Duplicates were removed using Scrublet^63^ (v0.2, python v3.6.8) by calculating the ‘doublet score’ for each barcode and removing barcodes with a ‘doublet score’ higher than the 90% quantile. One human sample (SN296) was removed during the cell type annotation procedure due to high contamination from neighbouring brain regions (see below). Altogether, 115,282 mouse, 180,956 human and 99,498 opossum cells, with a median of 2,392 UMIs per cell, passed the filtering steps (Extended Data Fig. 1a,d).

To assess the quality of the sequenced libraries, we aggregated expression values (mature mRNA counts) across all cells within each batch into pseudobulks and calculated Spearman’s rho correlation coefficients between the pseudobulks using genes expressed in at least 10% of the cells in at least one biological replicate (human n = 7,696; mouse n = 4,806; opossum n = 2,765). We observed high correlations between the libraries from the same developmental stage, even when different Chromium reagents (v2 and v3) were used to produce the libraries (Extended Data Fig. 1c).

### Data integration and clustering

We used LIGER (0.4.2, R)^14^ to perform batch correction and integrate data across stages within each species. Libraries across all developmental stages and individuals were considered as individual batches. Normalization and selection of highly variable genes were performed using LIGER with default parameters, followed by integrative non-negative matrix factorization (*optimizeALS* function) with k=100. The obtained embeddings were then used as the basis for UMAP visualization (uwot 0.1.10, R)^64^. To annotate cell types, we applied iterative unsupervised clustering within each species using the SCANPY (v1.5.1) implementation of the Louvain algorithm with a resolution of 3. First, we clustered the entire datasets for each species. For each identified cluster we split the data by batch and repeated the LIGER integration and Louvain clustering as described above to divide the data into subclusters. Batches with low cell contribution (<50 cells) to a cluster were excluded from sub-clustering. This iterative procedure yielded 68 (human), 61 (mouse) and 67 (opossum) clusters split into 574-611 subclusters for each species (Supplementary Tables 2-4, Extended Data Figure 2b).

We integrated the snRNA-seq data across species and across developmental stages in a common embedding (in pairs or all three species combined) using LIGER as described above. We used 1:1 orthologous genes detectable (i.e. at least 1 UMI) in all batches and variable across cells (n=6,101 for mouse/human; n=5,019 for mouse/opossum; n=3,742 for global). The initial integration resulted in manifolds where species-specific differences were still visible. To further merge the embeddings across species, MNN-correct^65^ (*fastMNN*, R, *batchelor package 1*.*0*.*1*) was applied with species assignment as the integration vector. This generated 100-dimensional aligned embeddings that were used for transfer of cell type labels (pairwise embeddings) or estimation of pseudoages (global embedding) as described below.

### Cell type annotation and label transfer

Given that the development of cerebellar cell types and their marker genes have been mostly described in mouse^6,12,7,66^, we first annotated the cells in the merged mouse dataset. Besides literature-based marker genes, we extensively used the *in situ* hybridisation data from the Allen Developing Mouse Brain Atlas^15^. We assigned each subcluster to a broad lineage (e.g. *VZ* for neurons born at ventricular zone), cell type (e.g., *Purkinje*) and cell state (e.g., *Purkinje_defined*). Cell states that displayed remaining variability were further split into subtypes (e.g., *Purkinje_defined_FOXP1*; Extended Data Fig. 2a-c). The cells that were not included in any subcluster were annotated only in case their identity could be unequivocally determined (Supplementary Table 2). The smallest clusters (<100 cells) were annotated as whole clusters. Although we mostly use the term “type” to group cells committed to a distinct mature cell fate and “state” to refer to differentiation status that often form a continuum within each cell type category (Extended Data Fig. 2a,c), there are a few notable exceptions: (1) splitting early neuroblasts based on their final fate was often not possible and we therefore included separate “cell type” level categories (e.g., *VZ_neuroblast*) to label these nascent postmitotic neurons; (2) the “cell type” level category *GC/UBC* was used to annotate subclusters that displayed co-expression of granule and unipolar brush cell markers; (3) we split neural progenitors into spatiotemporal “subtypes”, although some of these might alternatively be considered as sub-states of the same progenitor subtype (e.g., *progenitor_RL_early* and *progenitor_RL*); (4) in case of the *oligodendrocyte_progenitor* cell state the sub-states (*pre-OPC, OPC* and *COP*) were distinguished at the level of “subtype” categories; (5) we grouped all immune cells into a single “cell type” category, because of low numbers of these cells in our datasets. The terms used to designate cell state categories include descriptors “progenitor” (proliferating cells), “neuroblast” (nascent postmitotic neurons), “glioblast” (glial cells exiting cell cycle), “differentiating”, “defined”, “maturing”, “mature”, ordered from less differentiated to more mature cell states, but not formally aligned across different cell type lineages (Extended Data Fig. 2a,c). In sum, 97% of the cells in the mouse dataset were specified at the level of cell state, out of which 51% were additionally assigned to a subtype (Supplementary Table 2). 1.0% of cells were annotated only at the level of cell type, these included low-quality granule cells and unresolved subclusters that contain cells from different cell types located at the nuclear transitory zone (*NTZ_mixed*).

Next, we used the pairwise cross-species aligned embeddings to transfer the annotation labels from the mouse subclusters to the subclusters in the human and opossum datasets (Extended Data Figure 2b). We determined centroids of all species-specific subclusters in the aligned embeddings, calculated centroid Pearson correlations and transferred the label of the highest correlating mouse subcluster to each human and opossum subcluster. We curated the transferred labels by inspecting the expression of marker genes in each subcluster in the human and opossum datasets (Supplementary Tables 3-4). In the human dataset we noticed two clusters (23 and 34) that mostly contained cells from one batch (SN296) and where expression of *HOX* genes was detectable, indicating that these clusters contained contaminating brainstem cells. We removed this batch from the human dataset and did not annotate the remaining cells (n=113) in the two clusters. Out of the transferred subcluster labels, 69/61% (human/opossum) were confirmed, 4.2/9.9% were re-annotated because of sampling differences (e.g., midbrain-derived *MB_neuroblast* detected in the opossum dataset only), 23/19% of the labels were adapted at the level of subtypes or related cell states, 1.7/4.7% of the labels were changed at the level of cell type, and 2.8/4.8% of the labels were removed because of unclear identity of the cells in these subclusters (mixed populations of cells, likely doublets and/or low quality cells). The human and opossum subclusters that did not receive a transferred label, were annotated *de novo* as described for the mouse dataset.

As a result of the label transfer and curation procedures 98% and 94% of the cells in the human (not considering the removed brainstem library) and opossum datasets were specified at the level of cell state, out of which 47% and 40%, respectively, were additionally assigned to a subtype (Supplementary Table 3-4). 1.1% of human cells and 2.6% of opossum cells belonged to subclusters that contain cells from different cell types located at the nuclear transitory zone (*NTZ_mixed*). The unlabelled cells (2.5% of mouse, 1.2% of human and 3.7% of opossum cells) were included in the merged datasets, but excluded from downstream analyses of cerebellar cell type transcriptomes.

### Quantification of cell type abundances

To quantify the relative abundances of cell types at each developmental stage in mouse, human and opossum, we grouped cells by cell type and required at least 50 cells per group (this resulted in the removal of 0-280 cells per developmental stage). For Fig. 1d, we quantified the cell type proportions across the whole datasets. The human adult deep nuclei libraries were excluded from Fig. 1d, since we did not sample deep nuclei separately in the two other species. The cell type proportions in individual adult human samples from different cerebellar cortical regions and deep nuclei are shown in Extended Data Fig. 7c. For Fig. 1e, we quantified the cell type proportions in each biological replicate separately (human 8 wpc libraries from the same individual produced with the same Chromium version were merged; human adult deep nuclei libraries were excluded), and determined the median between the biological replicates. In Fig. 1e, we did not consider cells that were not assigned to a cell type or belong to cell types/subtypes, which are from brain regions adjacent to the cerebellum: GABA_MB, progenitor_MB, progenitor_isthmic, motor_neuron, neural_crest_progenitor, isthmic_neuroblast, MB_neuroblast.

### Bayes modelling of cell type abundance differences

To test differences in cell type abundances (Extended Data Fig. 7e), we applied a Bayesian hierarchical model to account for species-specific, biological and technical variability. Specifically, we modelled the number of cells of a certain cell type, *y*_*b*_ as follows:

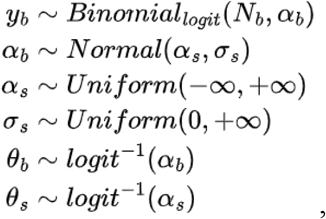

where *N*_*b*_ is the total cell count and α _*b*_ the logit transformed investigated cell type proportion within a distinct biological replicate. The hyperparameter α _*s*_ estimates the species-specific logit transformed abundance of a certain cell type. We apply the inverse logit to map α back to proportion scale (θ). This model was fitted using RStan^67^ (v2.19.3, R, *sampling* function; *iter=4,000, control=list(adapt_delta=0*.*99)*). To estimate the abundance disparity, we computed the pairwise differences of θ between the species, using the simulated posterior distribution and evaluating the 90% highest density interval (HDI^68^, *HDInterval*, v0.2.2, R) of the results. If the HDI did not overlap 0, we assumed a strong difference in cell type proportion.

### Identification of overdispersed genes

To stratify for informative genes, we identified highly variable genes based on an overdispersion cutoff. This approach was used for the estimation of correspondences between developmental stages across species (see *Stage correspondences across species*), comparing cell states and subtypes between datasets (see *Cross-species correlations and comparisons to the adult mouse data*), and identification of genes that are dynamic during neuronal differentiation (see *Gene expression trajectories*). Gene expression was normalized by dividing the number of UMIs by the sum of UMIs per cell (size factor). Next, we calculated the mean expression and variance using the normalized values to compute the gene-wise variance to mean ratio (*VMR*). Assuming Poisson distribution of non-informative genes, we estimated the expected variance to mean relationship (pVMR) by averaging over the inverse size factor. We expected genes that are highly variable to exhibit:

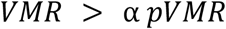

We set α, if not stated otherwise, to 1.5.

### Pseudoages

The 100-dimensional cross-species integrated global embedding of human, mouse and opossum datasets was used to determine a quasi-aligned continuous developmental vector, called pseudoage^18^. First, each mouse cell was assigned with a simple index of development, depending on its developmental stage, i.e. the first sampled stage (E10.5) was assigned with 1, the second (E11.5) with two. Next, this dataset was subsampled for each mouse time point to either 1,000 cells or the maximum number of cells per cell type, whichever value was smaller, to reduce the effect of cell type abundance differences between the stages. The resulting matrix was used as the reference. Next, for each cell in the human, mouse and opossum datasets, the 25 nearest reference cells (mouse) were identified and the mean of their developmental index was assigned to each query cell as its pseudoage.

### Stage correspondences across species

To establish correspondences between the developmental stages sampled in mouse, human and opossum, we applied three different metrics: (I) transcriptome similarity, (II) pseudoage similarity, and (III) cell state abundance agreement.

I. Transcriptome similarity. We identified the pairwise shared informative 1:1 orthologous genes in the human, mouse and opossum datasets (intersect of overdispersed genes, human vs. mouse n = 336, opossum vs. mouse n = 369 genes). We generated pseudobulks for each developmental stage by summing the UMI counts per gene from all cells. For each species, we normalized the expression values to CPM and mean-centered each gene’s expression values. Next, we subsetted the normalized data for the pairwise shared informative genes, to calculate the Spearman correlation coefficients between the developmental stage pseudobulks from all species, and computed the correlation distances by taking the arccosine of the distance.
II. Pseudoage similarity. We globally binned the cells from all species based on their pseudoages into 50 equally-sized bins. We then inferred for each developmental stage the proportions of cells in different pseudoage bins and used these proportions to calculate Manhattan distances for all cross-species developmental stage pairs.
III. Similarity of cell state proportions. For each cross-species pair of developmental stages, we computed the proportion of cells that had the same cell state annotation, and used these proportions to determine the pairwise Manhattan distances.

We used the dynamic time warping algorithm (dtw^69^, dtw v1.20, R) to find the lowest distance path through the 2D developmental plane in the distance matrices from all three approaches. We used mouse as a focal species for assigning correspondences. The three metrics showed good overall agreement (Extended Data Fig. 2d-f). When a developmental stage in human or opossum aligned with two or more stages in mouse, we kept the one with the smallest transcriptome correlation distance (Extended Data Fig. 2d-f). We grouped human samples from Carnegie stages 18 and 19 (7 wpc) as both show low transcriptome correlation distance to E11.5 in mouse. The toddler stage in human and P60 stage in opossum best matched P14 and/or P63 stages in mouse, but were kept as a separate intermediate stage between P14 and P63 in mouse, given that mouse P14 was assigned to the infant stage in human and P42 in opossum, and mouse P63 was assigned to the adult stage in both human and opossum.

### Gene expression scores

To quantify the expression of a group of genes of interest (GOI), we calculated gene expression scores akin to the approach used by La Manno *et al*.^18^. UMI data was normalised by calculating counts per million (CPM), and subsetted for the GOI. Next, we scaled the genes’ expression vectors to mean zero and unit variance, we averaged the scaled expression of all GOI to compute the score and calculated its 0.01 and 0.99 percentile. These percentiles were used for capping the score to eliminate outliers. Values outside this boundary were assigned to the nearest accepted value. This approach was used to quantify the expression of genes related to cell cycle^18^ (Extended Data Figs. 6g and 7a) or genes that gained expression in different cell types in the human lineage (Fig. 4h, Extended Data Fig. 10k).

### Cell type lineage assignments for ventricular zone neuroblasts

Although the VZ neuroblasts could not be separated into the associated terminal cell types (parabrachial and noradrenergic cells, GABAergic deep nuclei neurons, Purkinje cells, interneurons) based on clustering, we observed differential expression of known lineage-specific marker genes between developmental stages, in line with the known temporal order of the emergence of these cell types in the cerebellar ventricular zone^6,20^. We thus further split the mouse VZ neuroblasts based on their developmental stage: E10.5 cells were assigned to the parabrachial/noradrenergic cell type lineage, E11.5 cells to GABAergic deep nuclei neuron lineage, E12.5 and E13.5 to Purkinje cell lineage, and all other cells into interneuron lineage. In human, VZ_neuroblast_3 cells from Carnegie stage 18 were assigned to the parabrachial/noradrenergic cell type lineage, VZ_neuroblast_1 cells from Carnegie stages 18-19, VZ_neuroblast_2 cells from Carnegie stages 18-22 and VZ_neuroblast_3 cells from Carnegie stages 19-22 were assigned to the Purkinje cell lineage, and all other VZ neuroblasts to the interneuron lineage. In opossum, some VZ neuroblast subclusters could be assigned based on marker genes: cells in *LMX1A* and/or *LMX1B* positive subclusters (orig.cl_27_3, orig.cl_39_0, orig.cl_39_1, orig.cl_39_6) were assigned to the parabrachial/noradrenergic cell type lineage, and cells in *PAX2* and/or *SLC6A5* positive subclusters (orig.cl_23_9, orig.cl_26_10, orig.cl_31_2, orig.cl_31_4, orig.cl_31_5) to the interneuron lineage. The rest of opossum VZ neuroblasts were split based on their developmental stage: E14.5 and P1 cells were assigned to the GABAergic deep nuclei neuron lineage, P4 and P5 cells to Purkinje cell lineage, and P14 and P21 cells to the interneuron lineage. These assignments were used to visualize the VZ cell type lineages and to study the expression patterns of cell type marker genes amongst the VZ neuroblasts (Extended Data Fig. 3e,f), to perform LIGER integration for Purkinje and interneuron cell type lineages within each species (Extended Data Fig. 3g,j), and to integrate cells in Purkinje cell type lineage across species (Fig. 3d; see also *Pseudotemporal ordering*).

### Integration of data by subsets

For visualization purposes we integrated data by different subsets of cells belonging to a broad lineage, to a cell type lineage, or to a developmental stage. In this analysis, interneurons from adult human deep nuclei samples (n=57) were not included in the human interneuron subset, given that these were low in numbers, co-clustered with the molecular layer interneurons, and were not distinguished in adult mouse and opossum datasets, for which we only sampled whole cerebella. The integration of different subsets was performed as described for the whole datasets (see *Data integration and clustering*) using LIGER^14^. The number of components used for integrative non-negative matrix factorization (*optimizeALS* function) was as follows: Purkinje cell type lineage in mouse k=70, human k=50, opossum k=70 (Fig. 2a, Extended Data Fig. 3g); interneuron cell type lineage in mouse k=15, human k=20, opossum k=20 (Extended Data Fig. 3j); VZ broad lineage in mouse k=40, human k=50, opossum k=30 (Extended Data Fig. 3b,e); RL/NTZ broad lineage in mouse k=15, human k=20, opossum k=15 (Extended Data Fig. 4b,c); RL/EGL broad lineage in mouse k=30, human k=40, opossum k=40 (Extended Data Fig. 5b,c); glia broad lineage in mouse k=30, human k=40, opossum k=70 (Extended Data Fig. 6d). For the integration of individual developmental stages (Extended Data Figs. 3h, 7b) we used k=25.

### Cross-species correlations and comparisons to the adult mouse data

To match cell states and subtypes across species, we calculated Spearman correlation coefficients for all cross-species pairs within a subset (a broad lineage or a cell type). Interneurons from adult human deep nuclei (n=57) were excluded from this analysis (see also *Integration of data by subsets*). We used only the 1:1 orthologous genes that were overdispersed within a subset in each of the three species (i.e. intersect of highly variable genes). The number of the used genes for each subset is specified in the figure legends (Fig. 2d, Extended Data Figs. 3c,n, 4f,h, 5e,f, and 6e,f).

We used the same approach to match mouse Purkinje and granule cell developmental subtypes defined in this study to the adult subtypes reported by Kozareva *et al*.^7^, using all mouse genes that were overdispersed in both datasets (i.e. intersect of highly variable genes; Extended Data Figs. 3i,m, and 5j). We note that the adult mouse dataset^7^ was produced by methods similar to the ones used in this study: nuclei were extracted from frozen tissue samples dissected from different lobules of the cerebellar cortex and the libraries were prepared using v3 Chromium reagents (10x Genomics).

### Principal components analysis

To evaluate global relationships among the cell type-specific transcriptomes during development in the three species, we performed principal components analysis (PCA) using three-way 1:1 orthologous genes. Cells originating from the same biological replicate and assigned to the same cell type were merged into pseudobulks by summing up the UMI counts per gene. Only pseudobulks that contained at least 150 cells were considered. Only genes which were expressed in at least 10% of cells in a single pseudobulk in any species and showed variability in all species (variance(CPM) > 0) were kept for downstream analysis. The data was normalized to CPM and gene expression was median-centered within each species and aggregated. The first 15 principal components of the merged dataset were approximated using *prcomp_irlba* (irlba^70^ v2.3.3, R).

### Marker gene identification

To identify genes enriched in different cell states, term frequency - inverse document frequency (TF-IDF) transformation was applied (quickMarker function from the R package SoupX)^71^. *P*-values were computed using a hypergeometric test and corrected with the Benjamini Hochberg method. The count matrix was binarized with a threshold of > 0 UMI per cell. For each gene its expression within a specific group of cells was contrasted against all other cells in the dataset.

Conserved marker genes were called using an adjusted dataset. To reduce the effect of sampling differences between species, the data was aligned between the species: for each cell state, only those matched developmental stages were used, where the cell state is present in all three species. Next, each cell state per matched developmental stage was sampled to 1,000 cells, if needed the group was randomly upsampled. This dataset was then used for marker gene identification as described above. Conserved markers were defined as genes that are enriched in all species in the same cell state with a corrected *P*-value of less than 0.01, display cell state-specific enrichment of > 2 and show expression (UMI > 1) in at least 10% of the cells. Conserved markers were ranked by maximum scaling the TF-IDF value per species and calculating a score as follows:

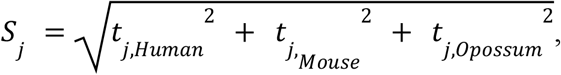

where *t* is the scaled TF-IDF value and *S* the ranking score per gene *j*.

#### Pseudotemporal ordering

We extracted from all three datasets the cells that were assigned to the Purkinje cell (Extended Data Fig. 3e,g) or granule cell (Extended Data Fig. 5b) lineage. We only used the three-way 1:1 orthologous genes that were detectable (number of UMIs > 0) in all species. We aligned the data from all species in the low dimensional space of 50 principal components using the Harmony pipeline^72^ (v1.0). Library and species identities were used as covariates. Next, we chose a starting cell, based on the UMAP embedding, and applied the SCANPY implementation of diffusion pseudotime^36^ (DPT) algorithm. Specifically, we used the Harmony-corrected principal components as input for the diffusion map algorithm and projected the cells into a ten-dimensional diffusion map, which together with the previously selected root cell was used as an input for the estimation of DPT values.

#### Gene expression trajectories

We used the cross-species aligned pseudotemporal cell orderings to model gene expression trajectories in all three species. First, we split the pseudotime vector into ten equally sized bins. For each bin, pseudobulks consisting of cells from the same biological replicate were merged by summing up the UMIs per gene, only considering the replicates where the number of cells was at least 50. Next, we determined for each pseudotime bin and species the mean UMI counts between the biological replicates. To identify the genes that are dynamic during neuronal differentiation, we filtered for highly variable genes (as described in *Identification of overdispersed genes*) using the pseudotime binned UMIs. In this case we set α in the highly variable gene formula (see above) to 1. Selection was done per species and the intersection of dynamic genes across species was used in the next steps. The count matrix was normalised, using all genes, to counts per million (CPM), subsetted for the highly variable genes and combined such that each orthologue was added as an individual feature, i.e. each gene appeared three times in the matrix, one time for each species. To infer groups of genes with similar trajectories, we used the soft clustering algorithm Mfuzz^73^ (v2.44.0) and clustered the genes into eight trajectory clusters. For each orthologue we determined its center of mass to infer its most prominent expression time window (Extended Data Figs. 9h, 10a). To order the trajectory clusters, we calculated the mean center of mass of all confident cluster members (cluster membership score > 0.5). The similarity of orthologues’ trajectories was tested by calculating a similarity *P*-value from the cluster membership score:

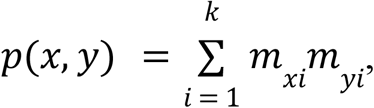

Where *m*_*xi*_ and *m*_*yi*_ is the cluster membership score for cluster *i* for the given orthologues from two species (*x* and *y*). The resulting *P*-value was adjusted for multiple testing using Benjamini-Hochberg method. Based on a set of rules, the orthologues were classified in multiple classes:

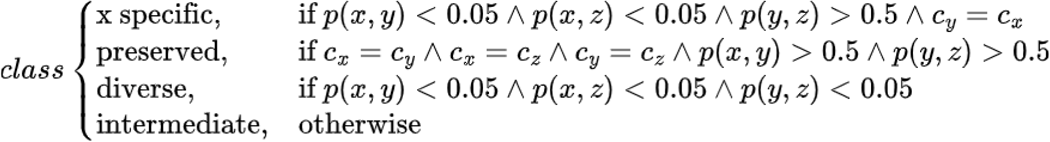

where *x, y, z* represent the studied species in any given combination, *p*(*x, y*) is the adjusted *P*-value for the comparison of the given orthologues in species *x* and *y, c*_x_ the cluster with the maximum membership score for the studied orthologue in species *x* (human-specific, mouse-specific, marsupial). If one or more of the three orthologues did not reach a maximum membership score above 0.5, the gene was excluded from downstream analysis. Orthologous genes with either species-specific or diverse trajectories were grouped into the ‘diverged’ class (Fig. 3g).

We additionally measured the similarities between the trajectories of the orthologous genes by computing dynamic time warping distances (dtw^69^ v1.20, R). Comparison of these distances for the groups of genes with either preserved, intermediate or diverged trajectories corroborated the clustering-based classification (Extended Data Fig. 9i). We further assessed the maximum and minimum pairwise dynamic time warping distances between the trajectories of the orthologues, to provide a quantitative measure of the amount of change for each gene (Figure 4c, Extended Data Fig. 10c).

The patterns of trajectory changes during Purkinje or granule cell differentiation were visualised using alluvial plots (ggalluvial^74^ v0.12.3, R). For both cell types, each orthologous gene was mapped to its trajectory cluster in each species, and coloured according to its trajectory cluster in human (Fig. 4a, Extended Data Fig. 10b).

### Presence/absence expression differences

To assess presence and absence of gene expression in cerebellar cell types, we generated pseudobulks by summing up exonic UMI counts (mature mRNA reference) per biological replicate, and cell type. For the meningeal and immune cell types, we summed up the counts from all cells in the dataset, because of their low abundance. We used the mature mRNA counts in this analysis, since most of the intronic reads arise due to internal priming from intronic polyA and polyT sequences^75^ that can differ between the species and thus the intronic counts cannot be directly compared. For each cell type, we only considered those matched developmental stages, for which there was data from all three species; and only those pseudobulks that had at least 50 cells. We normalized the data to CPM. In each species, we required a gene to reach a cutoff 50 CPM in at least 2 pseudobulks of a given cell type (or the single pseudobulk generated for meningeal or immune cells), to consider it as reliably expressed. For genes failing to meet this cutoff, i.e. not reliably expressed in any of the cell types, we also considered their expression in bulk RNA-seq RPKM-normalized data covering overlapping stages of cerebellum development in the same species^19^. Genes not detected as reliably expressed in the cerebellum (in any cell type) based on our snRNA-seq data but reaching a RPKM value above 5 in the bulk RNA-seq data, were assumed to be affected by technical artifacts of the snRNA-seq measurements. In these cases, we removed the orthologous gene group from the downstream analysis. 3,577 genes were affected by these technical limitations in at least one of the species and removed, whereas 9,189 1:1 orthologous genes were kept in the downstream analysis.

We then focused on the main glial and neuronal cell types in the cerebellum (astroglia, oligodendrocytes, GABAergic deep nuclei neurons, Purkinje cells, interneurons, glutamatergic deep nuclei neurons, granule cells, UBCs) to evaluate presence/absence of expression of three-way 1:1 orthologous genes. If a gene’s expression in two pseudobulks within a matched developmental stage reached the cutoff of 50 CPM, we assumed the gene to be reliably expressed in this cell type. Next, in each species we determined the maximum expression of each gene per cell type across all matched stages and calculated the pairwise fold differences between the orthologues. Each orthologous gene group was then classified in each cell type according to the following rules: (I) a gene is considered as *therian-expressed* if it is reliably expressed in all studied species; (II) a gene is considered as being expressed (present) in a species-specific manner, if it is reliably expressed in one species, but the maximum expression in the two other species is below the 50 CPM cutoff and the pairwise fold-differences are more than five-fold; (III) a gene is classified as not expressed (absent) in a species-specific manner, if its expression levels are below 50 CPM in one species, but it is reliably expressed in the other two and the pairwise fold-differences are more than five-fold. We then polarized the presence/absence expression differences between mouse and human, using opossum as an outgroup species, and assigned the genes as having gained or lost expression in the mouse or human lineage. The differences between the eutherian and marsupial lineages cannot be polarized, because of lack of a non-therian outgroup. Thus, if a gene is only expressed (present) in opossum, we classified it as *marsupial-expressed*. Similarly, if a gene is not expressed (absent) specifically in opossum, we classified it as *eutherian-expressed*.

### Gene expression cell type-specificity

To evaluate the cell type specificity of gene expression (Extended Data Fig. 10h), we determined specificity index τ, a metric described previously for evaluation of tissue-specificity^76^:

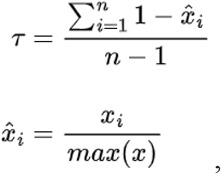

where *x* is the normalized expression value for a certain gene across multiple cell types within a given species; *x*_i_ is the cell type *i* specific expression value; *n* is the total number of observed cell types. We determined the τ separately for each species using all cell types that were considered in the analysis of presence/absence expression differences (astroglia, oligodendrocytes, GABAergic deep nuclei neurons, Purkinje cells, interneurons, glutamatergic deep nuclei neurons, granule cells, UBCs). The index value 0 indicates broad expression, and 1 cell type-specific expression.

### Gene ontology enrichment analyses

For gene ontology enrichment analyses considering three-way 1:1 orthologues, we extracted the mouse gene ontology data from ENSEMBL release 91. Genes were associated to all annotated GO terms and enrichments were calculated using a binomial test (*pbinom*, R) for the observed number of occurrences within a group, given the GO terms occurence in the full dataset. All *P*-values were adjusted for multiple testing with the Benjamini-Hochberg method.

### Adult bulk RNA-sequencing data analyses

The adult brain and cerebellum bulk RNA-sequencing data from nine mammals and chicken is from Brawand *et al*^*46*^. We used the reference genomes and orthology relationships from Ensembl release 91. Gene expression levels were measured as described by Wang *et al*.^60^. Briefly, for each gene we measured expression levels in the fragments per kilobase of coding DNA sequence (CDS) per million uniquely CDS-aligning reads (FPKM), a unit which corrects for both gene length and sequencing depth. We restricted the analysis to the coding regions of the longest protein-coding isoform of 1:1 orthologues that perfectly align across species.

### Essential and disease-associated genes

We used different metrics of gene essentiality: (i) LOEUF score (loss-of-function observed/expected upper bound fraction)^39^ from the Genome Aggregation Database (gnomAD), which ranks genes along a continuous spectrum of tolerance to loss-of-function variation based on human exome and genome sequencing data (*in vivo*); (ii) a measure of *in vivo* essentiality that combines various essentiality scores (RVIS, pLI, Phi, missense Z-score, LoFtool and s_het_) to rank genes based on human exome and genome sequencing data^38^; (iii) a measure of *in vitro* essentiality that combines viability scores from CRISPR-Cas9 screens on human cell lines^38^. We obtained the human inherited disease gene list from the manually curated Human Gene Mutation Database (HGMD, PRO 17.1)^42^, and only used genes with disease-causing mutations as described previously^41^. We used the genes that based on the mapping to the Unified Medical Language System (UMLS) were linked to the high level disease types ‘Nervous system’ or ‘Psychiatric’, and split these genes into two groups based on whether a gene was also linked to the high level disease type ‘Developmental’ (developmental n=373; other n=200). We obtained the cerebellum-linked disease gene lists from two sources. First, the curated lists of genes associated with neurodevelopmental and adult-onset neurodegenerative disorders linked to cerebellar dysfunction are from Aldinger *et al*.^13^. Briefly, the cerebellar malformation list includes 54 genes combining published Dandy–Walker malformation and cerebellar hypoplasia genes and genes identified through exome sequencing; the Joubert syndrome list includes 42 published Joubert syndrome genes; the autism spectrum disorder list includes 108 high-confidence genes identified through exome and genome sequencing; the intellectual disability list includes 186 genes identified through exome sequencing; the spinocerebellar ataxia list includes 44 genes associated with OMIM phenotype PS164400. Second, the lists of driver genes associated with paediatric cancers is based on Gröbner *et al*.^43^ and include 53 genes for medulloblastoma, 8 genes for ependymoma and 24 genes for pilocytic astrocytoma and pleomorphic xanthoastrocytoma.

Significance of categorical enrichments were evaluated using binomial tests using all dynamic genes as the background gene set (see *Identification of overdispersed genes* and *Gene expression trajectories*). Continuous differences were investigated using a permutation test (10,000 iterations) for *H*_1_ = *greater* (*in vivo / in vitro* essentiality scores) or *H*_1_ = *smaller* (LOEUF score). *P-*values were adjusted for multiple testing using Benjamini-Hochberg method.

### General statistics and plots

All statistical analyses and graphical representations were done in R (v.3.6.3) using the R packages: tidyverse^77^ v1.3.0; SingleCellExperiment^78^ v1.6.0; liger^14^ v0.4.6; rliger^79^ v1.0.0; batchelor^65^ v1.0.1; pheatmap^80^ v1.0.12; ggplot2^81^ v3.3.2. Additionally, the following Python (v3.6) packages were used: scanpy^82^ v1.5.1; htseq^83^ v0.13.5.

## Data and code availability

The datasets generated and analysed during the current study are available in the heiDATA repository, https://doi.org/10.11588/data/QDOC4E. Processed data can be interactively explored at https://apps.kaessmannlab.org/sc-cerebellum-transcriptome. Custom code is available at https://gitlab.com/kleiss/mammalian-cerebellum.

## Acknowledgements

We thank C. Conrad, A. Fallahshahroudi, F. Lamanna, D. Kawauchi, T. Trefzer and members of the Kaessmann group for discussions; M. Langlotz, T. Brüning, K. Mößinger, E. Renner, M. Toronyay-Kasztner, T. Nath Varma, B. Crespo Lopez, A. Billepp, P. Grimm and T. Wedig for assistance; J.L. VandeBerg for providing archived opossum samples; and the Joint MRC/Wellcome (MR/R006237/1) Human Developmental Biology Resource, Maryland Brain Collection at the Maryland Psychiatric Research Center (NIH NeuroBioBank), Chinese Brain Bank Center, and Human Brain Tissue Bank at Semmelweis University for providing human samples. The computational cluster bwForCluster of the Heidelberg University Computational Center is supported by the state of Baden-Württemberg through bwHPC and the German Research Foundation (INST 35/1134-1 FUGG). M.P. was supported by a grant from the Hungarian Brain Research Program (2017-1.2.1-NKP-2017-00002). M.C.-M. was supported by the Francis Crick Institute, which receives its core funding from Cancer Research UK (FC011171), the UK Medical Research Council (FC011171), and the Wellcome Trust (FC011171). This research was supported by grants from the European Research Council to H.K. (615253, OntoTransEvol) and S.M.P. (819894, BRAIN-MATCH).

## Author contributions

M.S., K.L., S.P. and H.K. conceived and organized the study; M.S., P.G., P.K., S.L. and M.P. collected samples; M.S., K.L. and F.M. established methods; M.S. performed experiments with support from N.M., C.S. and J.S.; K.L. performed data processing and analyses with contributions from M.S., I.S. and E.L., and input from F.M. and N.T.; M.S. and K.L. annotated and interpreted the data. K.L. developed the web application. K.O., P.Y., L.S. and L.M.K. provided feedback and discussions. I.S., S.A. and M.C.M. provided key scientific advice; S.M.P. and H.K. supervised the study and provided funding; M.S., K.L. and H.K. wrote the manuscript, with input from all authors.

## Competing interest declaration

The authors declare no competing interests.

## Additional information

Supplementary Information is available for this paper.

Correspondence and requests for materials should be addressed to H.K., S.M.P., M.S. and K.L..

**Extended Data Fig. 1.**
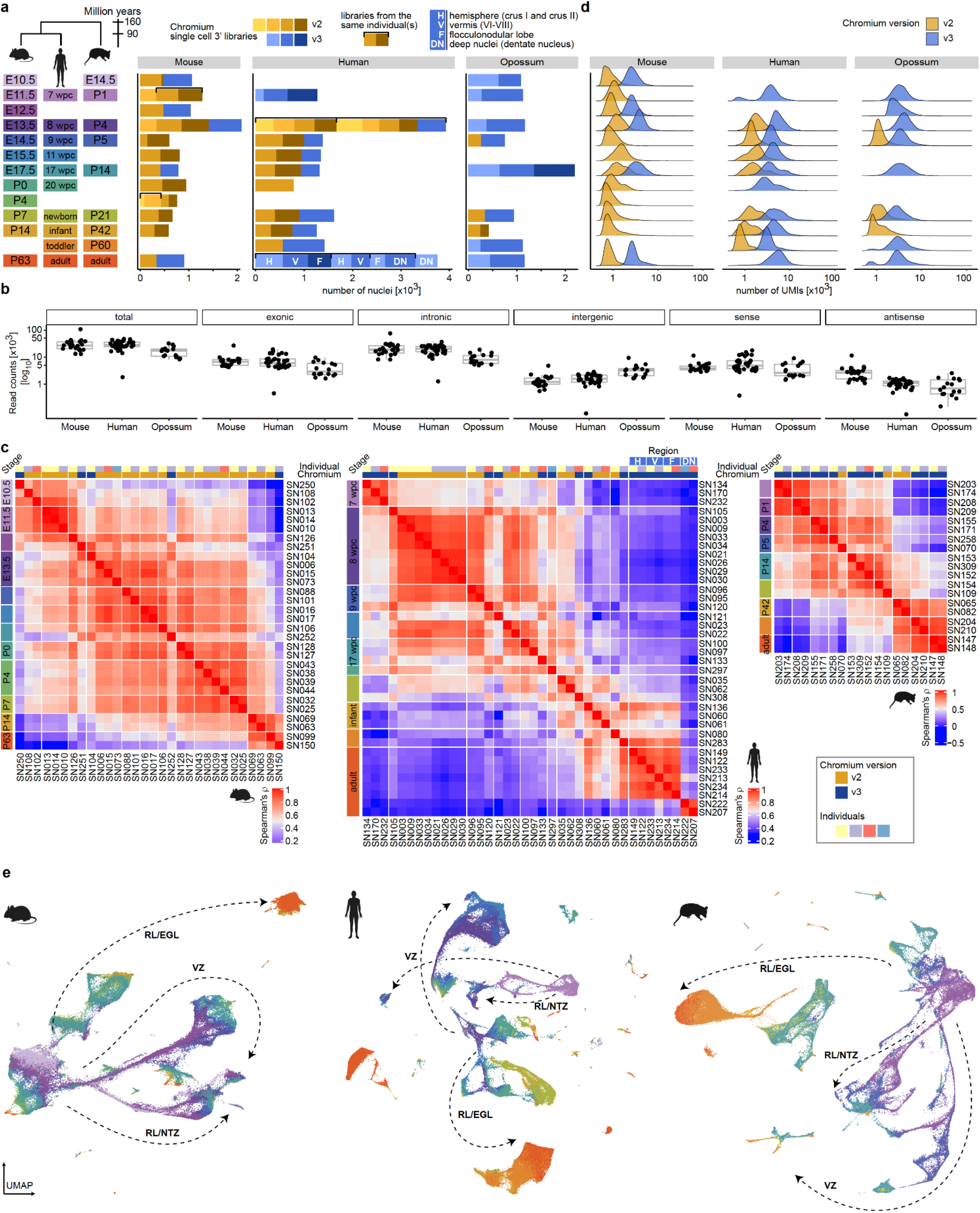
Overview of the datasets. **a**, Number of cells profiled by snRNA-seq per stage in mouse, human and opossum. A schematic of the sampled stages is shown on the left. Colours in bar plots indicate individual libraries (Chromium v2 in yellow hues and v3 in blue hues). Samples from different individuals were used for each library, except for the libraries grouped with brackets. The sampled cerebellum region is indicated for human adult libraries. **b**, Mapping statistics for the reads from mouse, human and opossum libraries. Shown are the total, exonic, intronic, and intergenic read counts per library based on the mature mRNA reference. Exonic counts are further split into sense and antisense read counts. Boxes represent the interquartile range, and whiskers extend to extreme values within 1.5 times the interquartile range from the box. **c**, Spearman’s rho correlation coefficients across libraries in mouse, human and opossum datasets. UMIs counted in mature mRNA mode were aggregated across all cells in each library. Correlations were calculated using the ranks of CPM (counts per million) values of the expressed genes (genes expressed in at least 10% of cells in any pseudobulk; human n = 7,696; mouse n = 4,806; opossum n = 2,765) within each library. Libraries are grouped by developmental stage and Chromium version. The sampled cerebellum region is indicated for human adult libraries, as in **a. d**, Distribution of UMI counts in v2 and v3 libraries in mouse, human and opossum datasets. **e**, Uniform Manifold Approximation and Projection (UMAP) of 115,282 mouse, 180,956 human and 99,498 opossum cells coloured by their developmental stage. Colours as in **a**. The broad neuronal lineages are shown with arrows. EGL, external granule cell layer; NTZ, nuclear transitory zone; RL, rhombic lip; VZ, ventricular zone.

**Extended Data Fig. 2.**
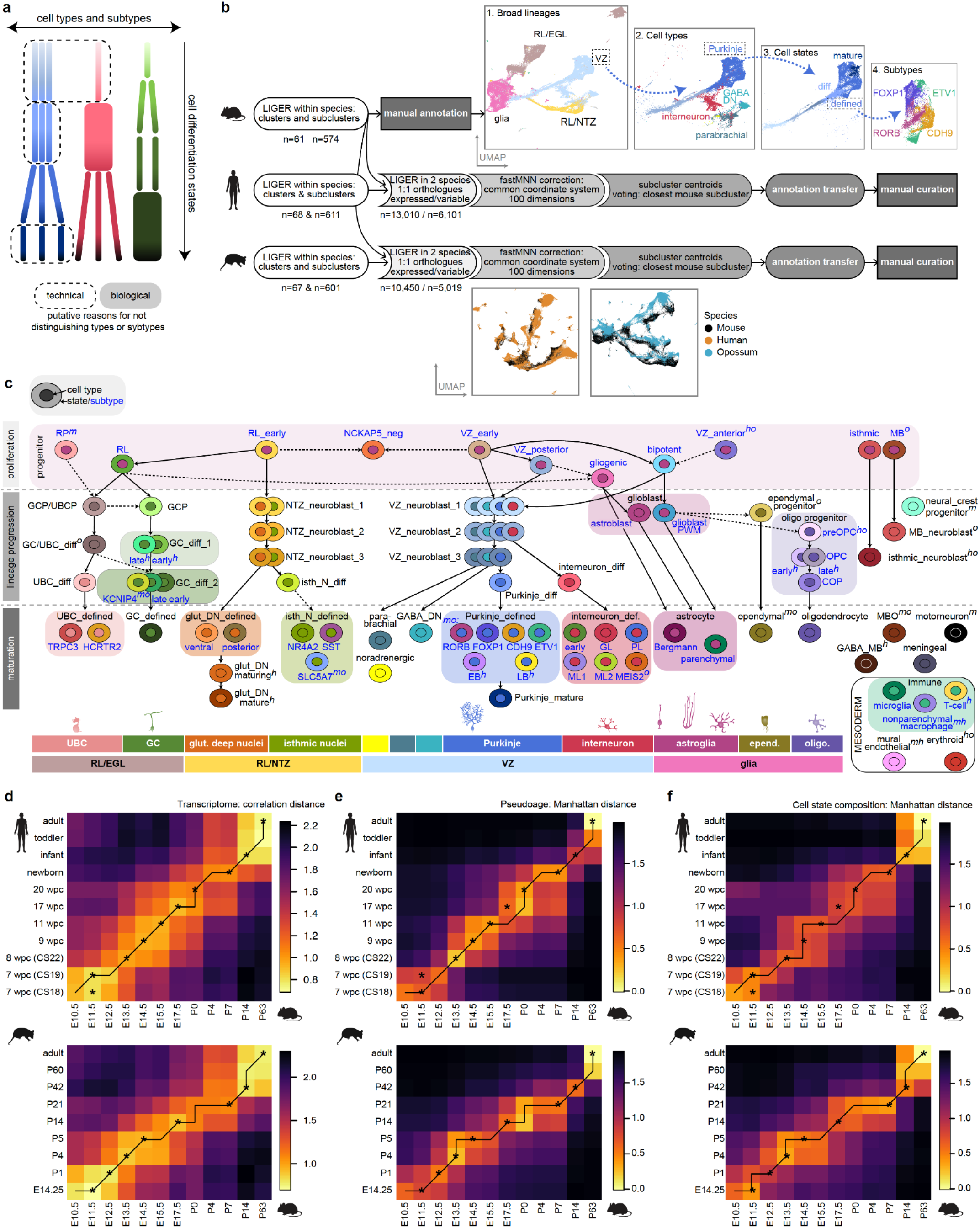
Cell type annotation and stage correspondences. **a**, Schematic summary of the annotation strategy. We used the term “type” to group cells committed to a distinct cell fate, and “state” to refer to differentiation status that often form a continuum within each cell type category. Shown are 3 hypothetical cell types (colous) and their subtypes (rectangles). Note that the cell state categories do not necessarily align across cell types. Both biological and technical reasons could explain why subtypes cannot be distinguished across all states in a given cell type. **b**, Schematic summary of the procedures used for the annotation of cell types in the mouse, human and opossum datasets. **c**, Overview of the cell type annotation categories. The schematic cells represent the different annotation categories with nucleus coloured by cell type and cytoplasm coloured by state or subtype. Cell state labels are in black, subtype labels in blue. Cell states at which subtypes were distinguished are highlighted with coloured background. For the categories not detected in all species, superscript text specifies the dataset(s) where a category is present: *h*, human; *m*, mouse; *o*, opossum. Solid arrows depict known lineage relationships, dashed lines depict additional relationships suggested by our data. Broad and cell type lineage groups are indicated at the bottom. **d-f**, Pairwise correspondences of developmental stages across species. The line indicates the best alignment between the time series determined by dynamic time warping algorithm using pseudobulk transcriptome correlation distances (**d**), Manhattan distances of pseudoages (**e**), and Manhattan distances of the cellular compositions at the level of cell states (**f**). Mouse was used as the focal species. Asterisks indicate the consensus stage correspondences from the three analyses. In **d**, we only used the genes that were informative in both datasets (intersect of overdispersed genes, human vs. mouse n = 336, opossum vs. mouse n = 369 genes). COP, committed oligodendrocyte precursor; def., defined; diff, differentiating; CS, Carnegie stage; DN, deep nuclei; EB, early-born; GABA, GABAergic; GC, granule cell; GCP, granule cell progenitor; glut, glutamatergic; isth N, isthmic nuclei neurons; LB, late-born; MB, midbrain; MBO, midbrain-originating cell; NTZ, nuclear transitory zone; oligo, oligodendrocyte; OPC, oligodendrocyte progenitor cell; preOPC, precursor of oligodendrocyte progenitor cell; PWM, prospective white matter; RL, rhombic lip; RP, roof plate; UBC, unipolar brush cell; UBCP, unipolar brush cell progenitor; VZ, ventricular zone; wpc, weeks post conception.

**Extended Data Fig. 3.**
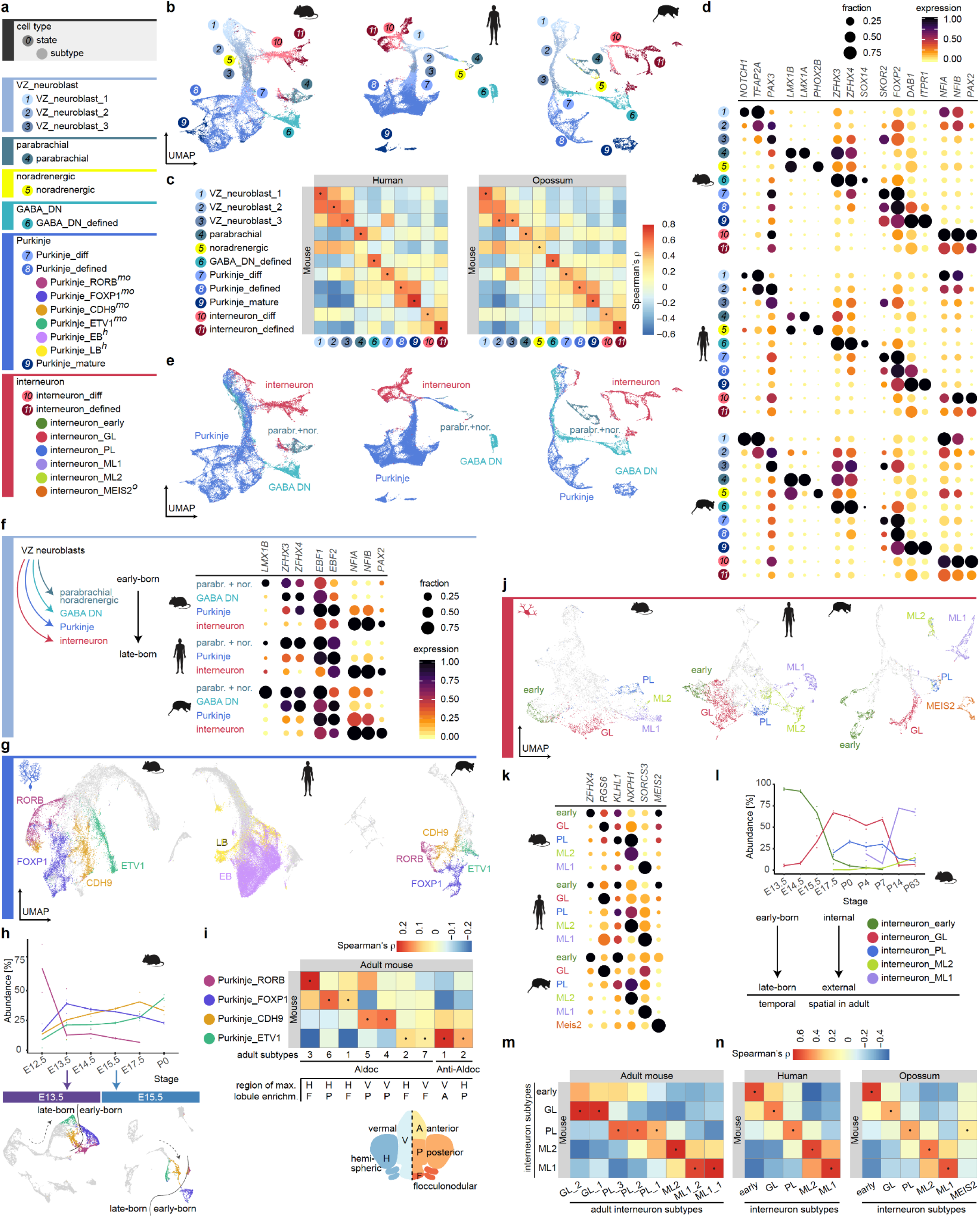
Atlas of the VZ cell types. **a**, Cell types, states and subtypes of neurons born at the ventricular zone. For the categories not detected in all species, superscript text specifies the dataset(s) where a category is present: *h*, human; *m*, mouse; *o*, opossum. **b**, Uniform Manifold Approximation and Projection (UMAP) of 37,391 mouse, 61,585 human and 22,674 opossum VZ-derived cells coloured by their state. Colours and numbers as in **a. c**, Spearman’s correlation coefficients between orthologous variable gene (n=208) expression profiles from mouse, human and opossum VZ-associated cell states. **d**, Expression of key marker genes in the VZ-associated cell states in mouse, human and opossum. **e**, UMAPs as in **b** coloured by cell type lineages. VZ neuroblasts were split into lineages giving rise to the different mature cell types based on the information about their developmental stage and marker gene expression. **f**, Expression of key marker genes in the VZ neuroblasts split into lineages as in **d** in mouse, human and opossum. **g**, UMAPs of 23,255 mouse, 49,399 human and 8,973 opossum Purkinje lineage cells coloured by their subtype. Subtype colours as in **a**; neuroblasts, early differentiating and mature Purkinje cells are in grey. **h**, Top: Purkinje subtype relative abundances (median of biological replicates) across developmental stages in mouse. Bottom: UMAPs of all mouse E13.5 and E15.5 cells, Purkinje subtypes are highlighted with colours. The dashed arrow directs from less mature cells (VZ neuroblasts) to more mature cells (defined Purkinje cells). The line separates early- and late-born Purkinje subtypes. **i**, Spearman’s correlation coefficients between shared variable gene (n=337) expression profiles from mouse Purkinje subtypes from this study and adult subtypes described by Kozareva et al.^7^. For each adult subtype the position of the lobule showing the highest enrichment^7^ along the mediolateral and anteroposterior axes is indicated. **j**, UMAPs of 6,422 mouse, 7,640 human and 5,815 opossum GABAergic interneurons coloured by their subtype. Subtype colours as in **a**; neuroblasts and differentiating interneurons are in grey. **k**, Expression of key marker genes in the interneuron subtypes in mouse, human and opossum. **l**, Interneuron subtype relative abundances (median of biological replicates) across developmental stages in mouse. The temporal order of interneuron subtype emergence gives rise to the spatial order in the adult cerebellum. **m**, Spearman’s correlation coefficients between shared variable gene (n=329) expression profiles from mouse interneuron subtypes from this study and adult subtypes described by Kozareva et al.^7^. **n**, Spearman’s correlation coefficients between orthologous variable gene (n=198) expression profiles from mouse, human and opossum interneuron subtypes. In **c, i, m** and **n**, dots indicate the highest correlation for each column. In **d, f** and **k**, dot size and colour indicate the fraction of cells expressing each gene and the mean expression level scaled per species and gene, respectively. diff, differentiating; EB, early-born; GABA DN, GABAergic deep nuclei neurons; GL, granule cell layer; LB, late-born; ML, molecular layer; parabr.+nor., parabrachial and noradrenergic cells; PL, Purkinje cell layer.

**Extended Data Fig. 4.**
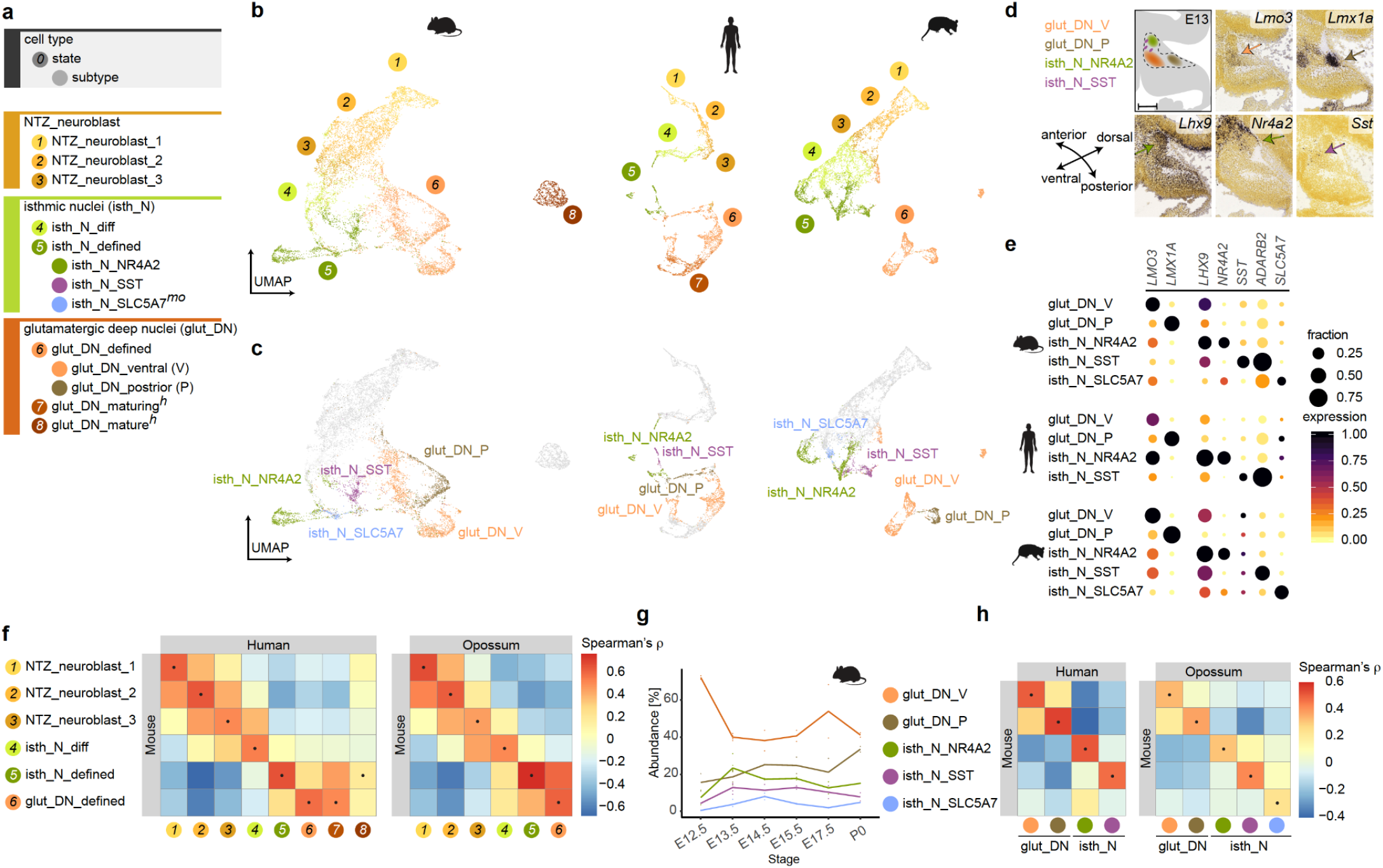
Atlas of the RL/NTZ cell types. **a**, Cell types, states and subtypes of neurons born at the early rhombic lip and/or located at the nuclear transitory zone during development. For the categories not detected in all species, superscript text specifies the dataset(s) where a category is present: *h*, human; *m*, mouse; *o*, opossum. **b, c**, Uniform Manifold Approximation and Projection (UMAP) of 10,949 mouse, 6,301 human and 9,965 opossum RL/NTZ cells coloured by their state (**b**) or subtype (**c**). Colours and numbers as in **a. d**, Spatial distribution of glutamatergic deep nuclei and isthmic nuclei subtypes in mouse E13.5 cerebellar primordium based on RNA *in situ* hybridisation data^15^ for subtype marker genes. Sagittal sections counterstained with HP Yellow are shown. A schematic summary is shown in the top left panel. Scale bar: 200 μm. **e**, Expression of key marker genes in the glutamatergic deep nuclei and isthmic nuclei subtypes in mouse, human and opossum. Dot size and colour indicate the fraction of cells expressing each gene and the mean expression level scaled per species and gene, respectively. **f, h**, Spearman’s correlation coefficients between orthologous variable gene expression profiles from mouse, human and opossum cell states (**f**; n=228 genes) or subtypes (**h**; n=225 genes) in the RL/NTZ broad lineage. Dots indicate the highest correlation for each column. **g**, Subtype relative abundances (median of biological replicates) across developmental stages in mouse. glut DN, glutamatergic deep nuclei neurons; isth N, isthmic nuclei neurons; P, posterior; V, ventral.

**Extended Data Fig. 5.**
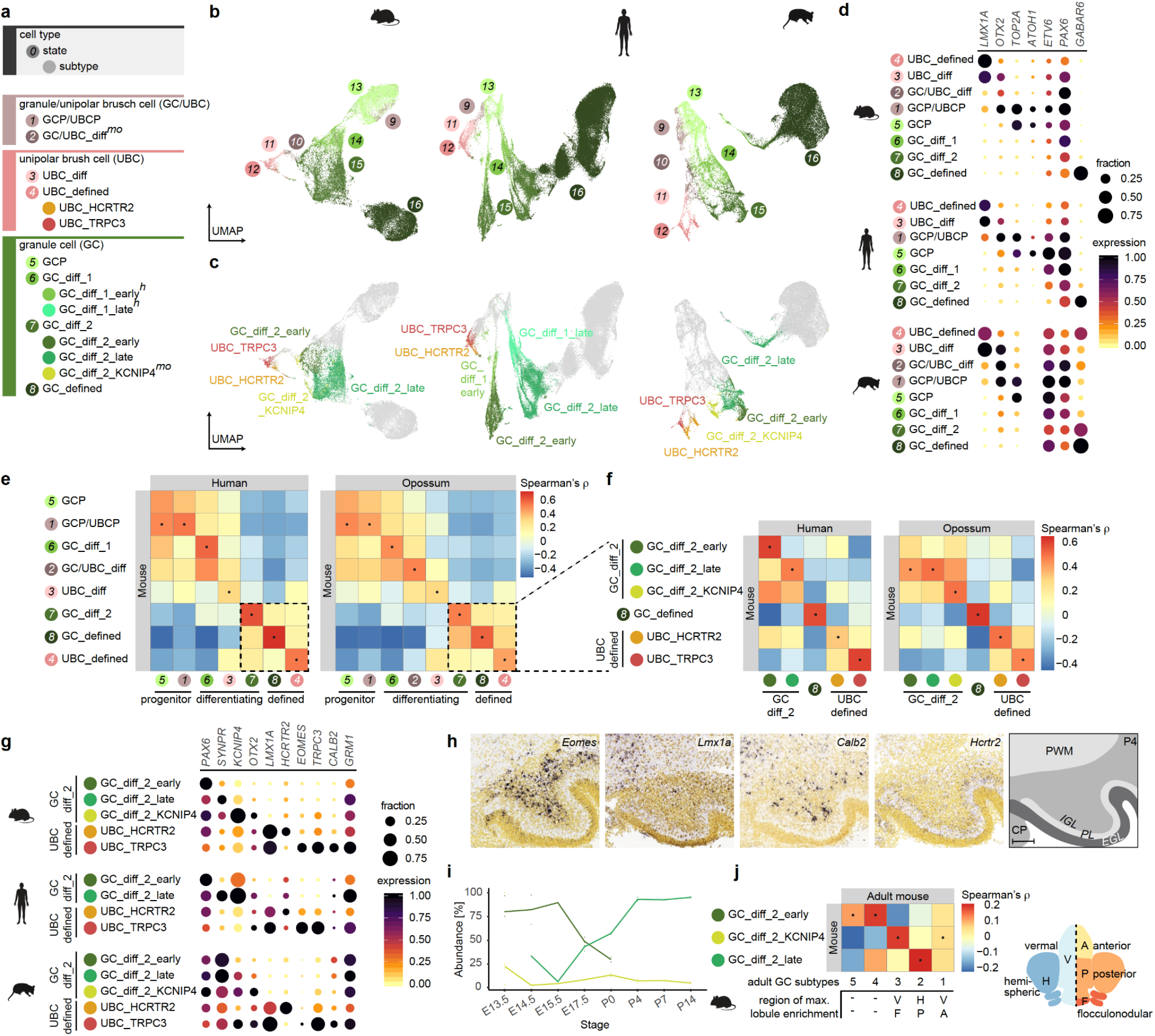
Atlas of the RL/EGL cell types. **a**, Cell types, states and subtypes of neurons born at the late rhombic lip associated with the external granule cell layer. For the categories not detected in all species, superscript text specifies the dataset(s) where a category is present: *h*, human; *m*, mouse; *o*, opossum. **b, c**, Uniform Manifold Approximation and Projection (UMAP) of 32,767 mouse, 73,492 human and 36,585 opossum RL/EGL cells coloured by their state (**b**) or subtype (**c**). Colours and numbers as in **a. d, g**, Expression of key marker genes in the granule and unipolar brush cell states (**d**) and subtypes (**g**) in mouse, human and opossum. Dot size and colour indicate the fraction of cells expressing each gene and the mean expression level scaled per species and gene, respectively. **e, f**, Spearman’s correlation coefficients between orthologous variable gene expression profiles from mouse, human and opossum cell states (**e**; n=110 genes) or subtypes (**f**; n=101 genes) in the RL/EGL broad lineage. **h**, RNA *in situ* hybridisation data^15^ for unipolar brush cell marker genes. An area of lobule X from mouse P4 sagittal sections counterstained with HP Yellow is shown for each marker gene. A scheme of layers in the P4 cerebellum is on the right. Scale bar: 100 μm. **i**, Relative abundances (median of biological replicates) of differentiating granule cell subtypes across developmental stages in mouse. **j**, Spearman’s correlation coefficients between shared variable gene (n=98) expression profiles from mouse differentiating granule cell subtypes from this study and adult subtypes described by Kozareva et al.^7^. For each adult subtype the position of the lobule showing the highest enrichment^7^ along the mediolateral and anteroposterior axes is indicated. In **e, f** and **j**, dots indicate the highest correlation for each column. CP, choroid plexus; diff, differentiating; EGL, external granule cell layer; GC, granule cell; GCP, granule cell progenitor; IGL, internal granule cell layer; PL, Purkinje cell layer; PWM, prospective white matter; UBC, unipolar brush cell; UBCP, unipolar brush cell progenitor.

**Extended Data Fig. 6.**
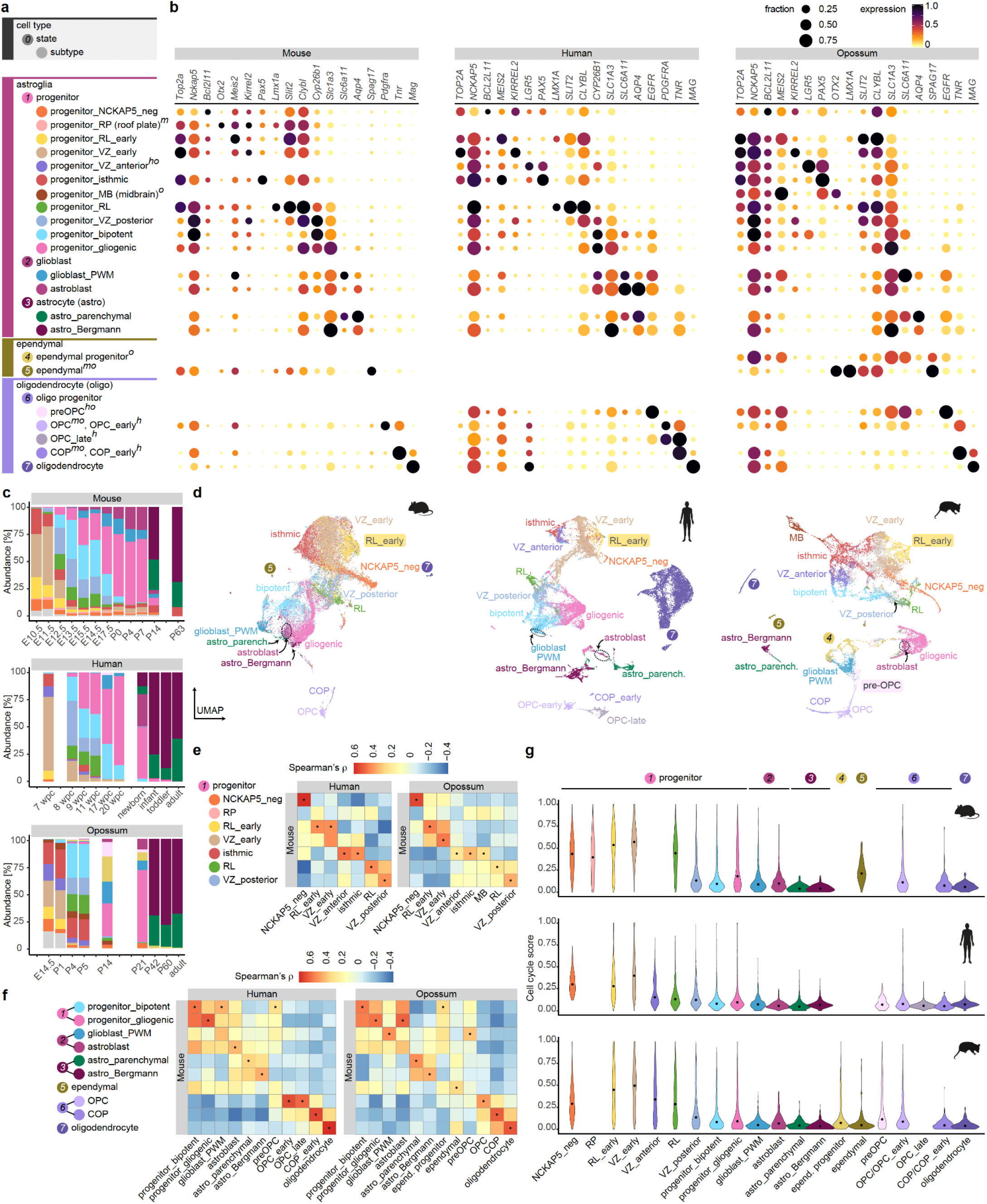
Atlas of the glial cell types. **a**, Cell types, states and subtypes of glial cells, including neural progenitor cells. For the categories not detected in all species, superscript text specifies the dataset(s) where a category is present: *h*, human; *m*, mouse; *o*, opossum. **b**, Expression of key marker genes in glial cell states and subtypes in mouse, human and opossum. Dot size and colour indicate the fraction of cells expressing each gene and the mean expression level scaled per species and gene, respectively. **c**, Relative abundances of astroglia subtypes, ependymal progenitors and preOPCs across developmental stages. Colours are as in **a**; astroglial cells not assigned to a subtype are in grey. Stages are aligned as in Fig. 1a. **d**, Uniform Manifold Approximation and Projection (UMAP) of 28,486 mouse, 32,897 human and 20,742 opossum glial cells coloured by their subtype or state. Colours and numbers as in **a**. Progenitors not assigned to a subtype are in grey. Mouse roof plate progenitors and human preOPCs are low in numbers and not discernible in this UMAP. **e, f**, Spearman’s correlation coefficients between orthologous variable gene expression profiles from mouse, human and opossum early progenitors (**e**; n=92 genes) or late progenitors and other glial cells (**f**; n=129 genes). Dots indicate the highest correlation for each column. **g**, Distribution of cell cycle score values across glial categories in mouse, human and opossum. Points indicate median score value. astro, astrocyte; COP, committed oligodendrocyte precursor; MB, midbrain; OPC, oligodendrocyte progenitor cell; preOPC, precursor of oligodendrocyte progenitor; PWM, prospective white matter; RL, rhombic lip; RP, roof plate; VZ, ventricular zone.

**Extended Data Fig. 7.**
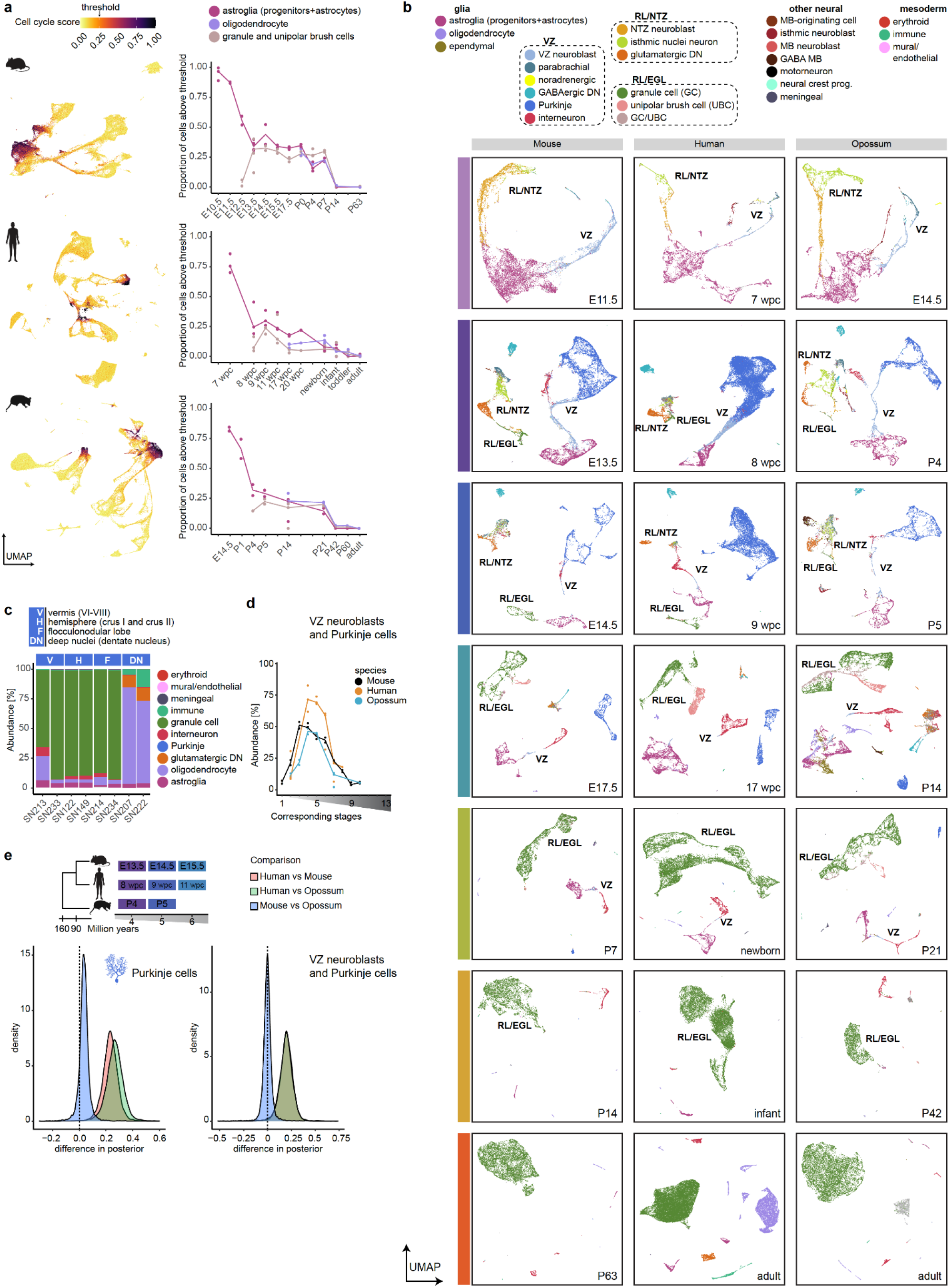
Dynamics of cell type abundances across development. **a**, UMAPs of mouse, human and opossum cells coloured by cell cycle score (left) and the proportion of cells (median across biological replicates) above the threshold score value (0.25) among astroglia, oligodendrocytes, and granule and unipolar brush cells (right). The stages are aligned as shown in Fig. 1a. **b**, Individual developmental stage UMAPs of mouse, human and opossum cells coloured by their cell type. Only the stages with correspondences in all studied species are shown. Labels indicate the broad neuronal lineages. **c**, Relative cell type abundances in individual adult human samples from different regions of the cerebellum. **d**, Relative abundances of cells annotated as Purkinje cells or VZ neuroblasts across developmental stages. Stages are aligned as in Fig.1a and the line indicates the median of biological replicates. **e**, Hierarchical Bayes model analysis of differences in the cell type abundances across species at corresponding developmental stages 4-6. Difference in posterior shows proportion differences of the fitted model. Vertical line indicates no shift in proportions. The modelled difference in Purkinje cell abundances can be summarised using a 90% highest density interval (HDI_90_) as follows; human vs mouse: [0.14, 0.32]; human vs opossum: [0.17, 0.37]; mouse vs opossum: [-0.023, 0.09]. To exclude the effect of possible biases in the annotation of VZ neuroblasts and Purkinje cells between the three species, both cell types were analysed together and the hierarchical model refitted. HDI_90_ of the posterior abundance differences: human vs. mouse: [0.09, 0.3]; human vs. opossum: [0.09, 0.31]; mouse vs. opossum: [-0.7,0.6].

**Extended Data Fig. 8.**
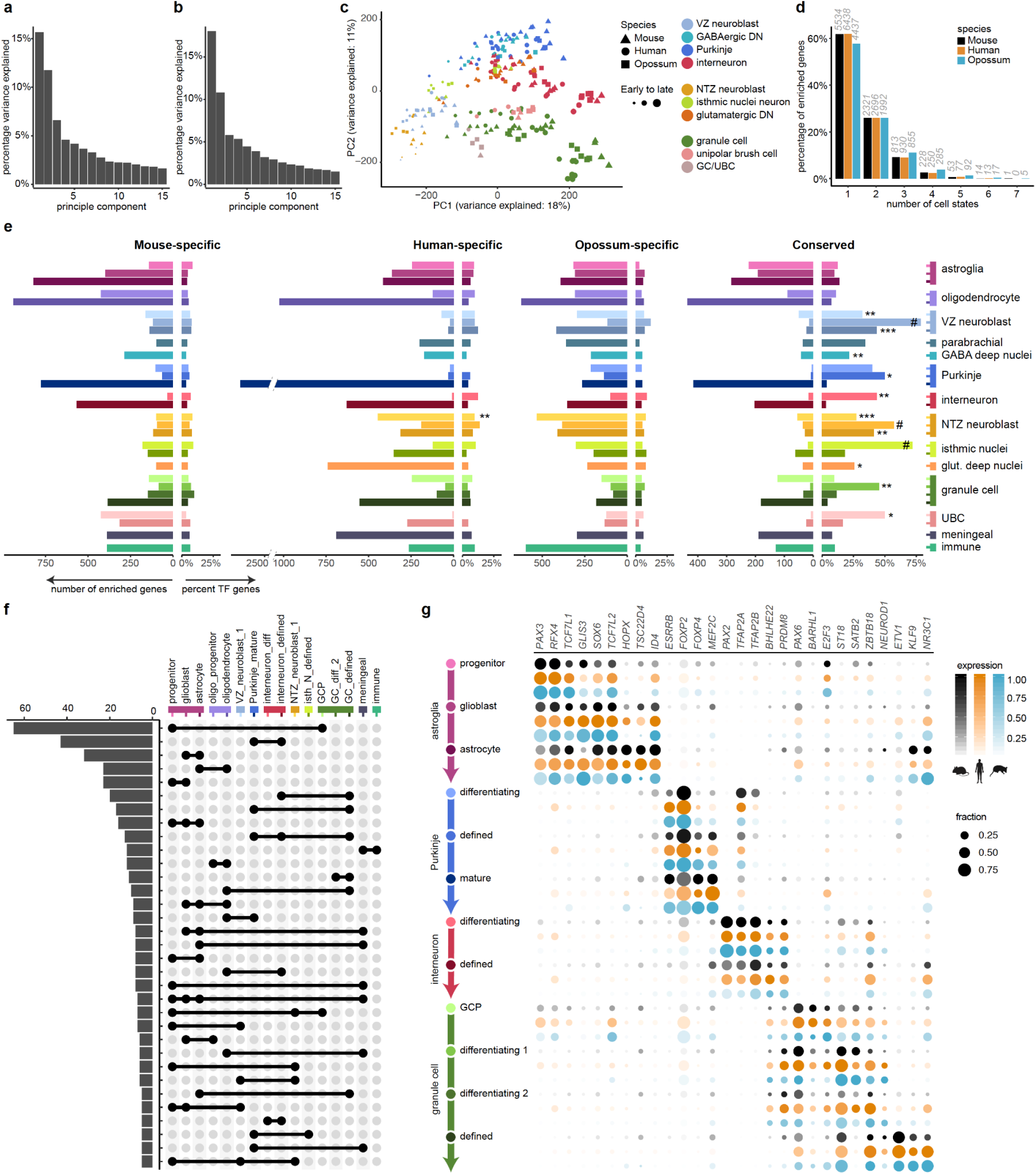
Transcriptional programs and marker genes. **a, b**, The percentage of variance explained by the first 15 principal components in the global PCA (**a**) and in the PCA of neurons only (**b**). **c**, PCA of neuronal cells based on 10,276 expressed orthologous genes across the three species. Data points represent cell type pseudobulks for each biological replicate. **d**, Number of cell states in which genes show expression enrichment in mouse, human and opossum. **e**, Numbers of markers and percentage of TF genes per cell state. Species-specific and conserved markers are shown. TF enrichments were identified using hypergeometric tests against a background of all 1:1:1 orthologous genes detected in the cell states included in the analysis; P-values were adjusted for multiple testing using Benjamini-Hochberg method; * *P* < 0.05, ** *P* < 10^−2^, *** *P* < 10^−3, #^ *P* < 10^−6^. **f**, Conserved markers shared between cell states. All groups with at least 5 genes are shown. **g**, Expression of TF genes that are among conserved markers across cell states of selected cell types in mouse (black), human (orange) and opossum (blue). Dot size and colour intensity indicate the fraction of cells expressing each gene and the mean expression level scaled per species and gene, respectively.

**Extended Data Fig. 9.**
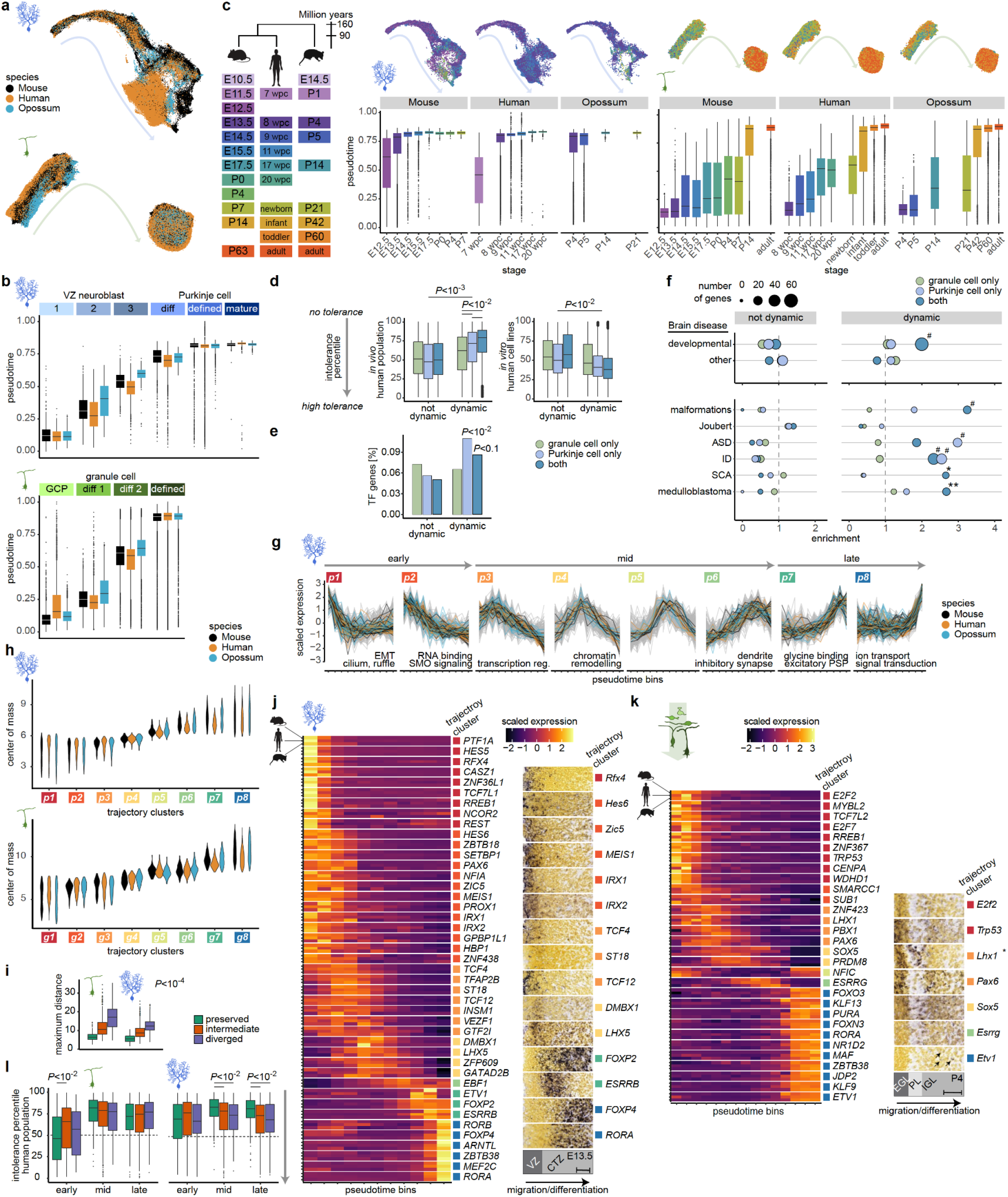
Gene expression trajectories across neuronal differentiation. **a**, UMAP of mouse, human and opossum cells assigned to the Purkinje cell (top) or granule cell (bottom) lineage and integrated across species. Cells are coloured by species. **b**, Pseudotime values across cell state categories for Purkinje (top) and granule (bottom) cell lineage. **c**, Pseudotime values across developmental stages in the mouse, human and opossum datasets for Purkinje (left) and granule (right) cell lineage. Stage correspondences are shown on the left and the integrated UMAPs plotted per species and coloured by stages at the top. **d**, Intolerance to functional mutations in human population (*in vivo*; left) or human cell lines (*in vitro*; right) for genes dynamic or non-dynamic during differentiation of granule cells, Purkinje cells, or both neuron types in all species. The integrated essentiality scores and distribution of human genes into percentiles as described by Bartha *et al*.^38^ **e**, Percentage of transcription factor genes across gene sets as in **d**. Adjusted *P*-values, binomial test. **f**. Enrichment of disease-associated genes across gene sets as in **d**. Top: inherited brain disease genes were split into two groups based on overlap with developmental disease genes^42^. Bottom: neurodevelopmental and neurodegenerative diseases^13^, and malignancies^43^ directly linked to cerebellar function and cell types. Adjusted \**P* < 0.1, \**P* < 0.01, ^#^*P* < 10^−4^, binomial test. **g**, Clusters of gene expression trajectories across Purkinje cell differentiation. Clusters (p1-p8) are ordered from early to late differentiation based on the mean center-of-mass values of the confident cluster members’ trajectories. Strongly preserved trajectories of the orthologues are highlighted with colours. Examples of enriched gene ontology categories for the genes with preserved trajectories are indicated. **h**, Center-of-mass values of individual trajectories across Purkinje (top) and granule (bottom) cell trajectory clusters. **i**, Maximum dynamic time warping aligned distance between the orthologues belonging to different trajectory conservation groups. **j, k**, Expression transcription factor genes with strongly preserved trajectories across Purkinje (**j**) and granule cell (**k**) differentiation. Scaled expression across pseudotime bins is shown on the left and RNA *in situ* hybridisation data^15^ on the right. Areas from sagittal sections of mouse E13.5 cerebellar primordia (**j**) or P4 cerebellar cortex in lobule III (**k**) counterstained with HP Yellow are shown. Schemes of layers in the respective areas are at the bottom. Arrows in **k** point to rare positive cells. \**Lhx1* is additionally expressed in Purkinje cells. Scale bars: 50 μm. **l**, Intolerance to functional mutations in human population (*in vivo*) for genes dynamic or non-dynamic during differentiation of granule cells, Purkinje cells, or both neuron types in all species. In **b, c, d, i** and **l**, boxes represent the interquartile range, and whiskers extend to extreme values within 1.5 times the interquartile range from the box. In **d, i** and **l**, adjusted *P*-values were calculated via a permutation test of pairwise comparisons between all categories (**d, i**) or trajectory conservation groups (**l**). ASD, autism spectrum disorders; CTZ, cortical transitory zone; diff, differentiating; EGL, external granule cell layer; ID, intellectual disability; IGL, internal granule cell layer; PL, Purkinje cell layer; PSP, postsynaptic potential; SCA, spinocerebellar ataxia; SMO, Smoothened receptor; VZ, ventricular zone.

**Extended Data Fig. 10.**
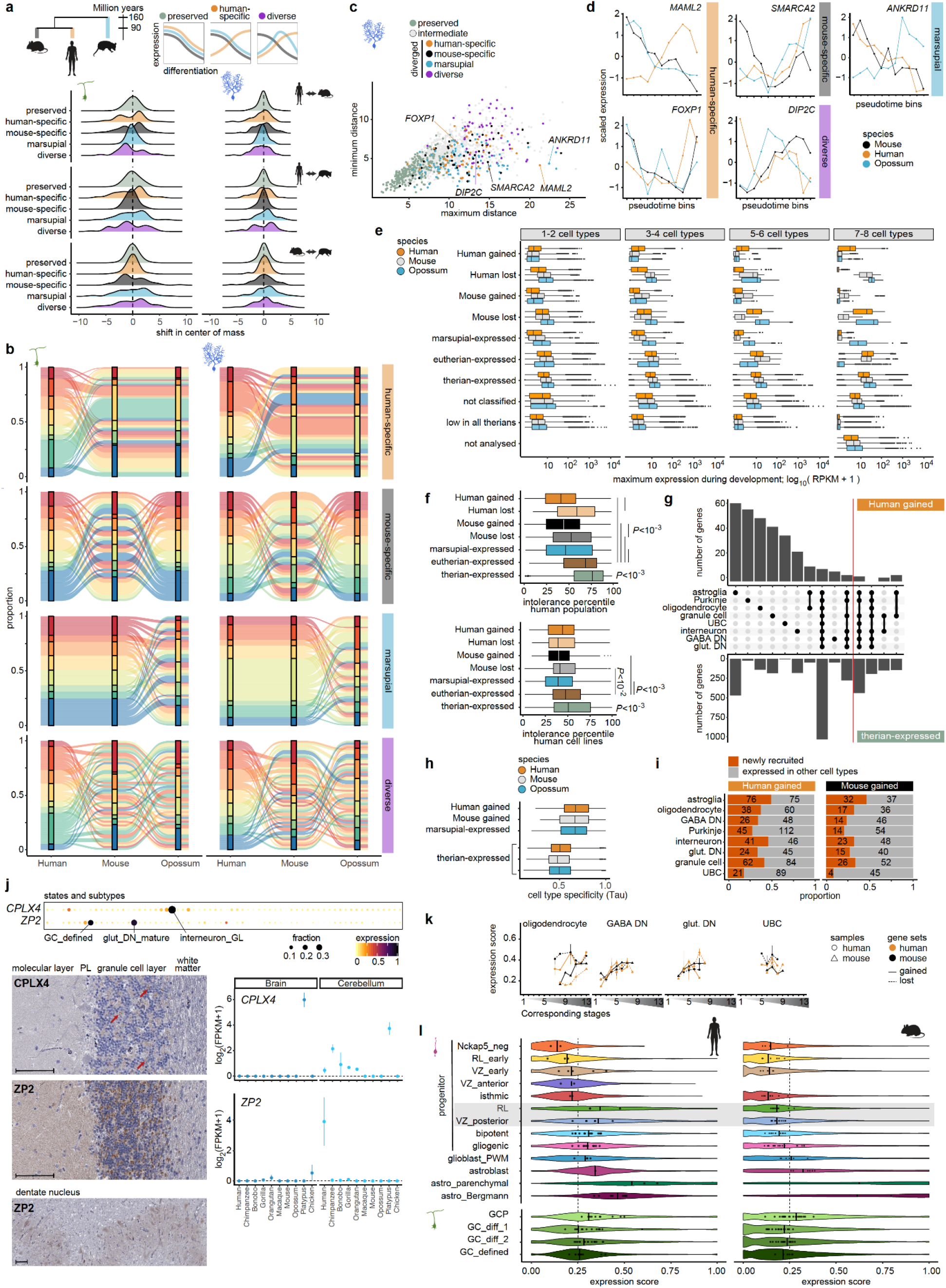
Differences in gene expression. **a**, Distribution of the pairwise (human vs mouse, human vs opossum, mouse vs opossum) shifts in the center of mass values of the trajectories that were assigned to preserved, human-specific, mouse-specific, marsupial and diverse patterns groups. The scheme (top) shows examples of a preserved pattern, a change in the human lineage, and a diverse pattern that cannot be assigned to a lineage. The changes in genes that differ between eutherians and opossum (marsupial) cannot be polarized. **b**, Distribution between trajectory clusters for genes with changes in gene expression trajectories during granule cell (left) or Purkinje cell (right) differentiation. **c**, Scatter plot of minimum and maximum pairwise distances between the expression trajectories of orthologues from the three species in Purkinje cell differentiation. High maximum and low minimum distances indicate the strongest lineage-specific trajectory changes. **d**, Examples of genes that display cross-species differences in trajectories during Purkinje cell differentiation. **e**, Comparison of cerebellar gene expression patterns in the snRNA-seq datasets to the bulk RNA-seq data^19^. Genes were assigned to groups based on the presence/absence expression patterns in the cerebellar cell types in the three therian species. Genes below the threshold (< 50 CPM) in all species are marked as low in all therians; for each species the genes that were below the threshold (< 50 CPM) in all cerebellar cell types based on snRNA-seq data, but were robustly expressed in the bulk RNA-seq data (> 5 RPKM), were excluded from the analysis (not analysed; n=3,577). Maximum expression across development, based on the bulk RNA-seq data, is shown. **f**, Intolerance to functional mutations in human population (top) or human cell lines (bottom) for genes assigned to different groups based on presence/absence expression pattern in the three species. Data from all cell types is summarised. The integrated essentiality scores and distribution of human genes into percentiles as described by Bartha *et al*.^38^ Adjusted *P*-values were calculated via a permutation test of pairwise comparisons. **g**, Number of genes assigned as human-gained or therian-expressed in different cerebellar cell types and their combinations. Ten cell type groups with the highest number of genes in the human-gained gene set (left from the red line) or therian-expressed gene set are shown. **h**, Cell type-specificity index (Tau, τ) for the genes that are robustly expressed in a single species or all species. Data from all cell types is summarised. The index ranges from 0 (broad expression) to 1 (restricted expression). **i**, Number of human-gained and mouse-gained genes that were expressed in other neural cell types in the cerebellum or newly recruited to the cerebellar transcriptome. **j**, Expression of ZP2 and CPLX4 in the human snRNA-seq dataset from this study (top), in human cerebellar sections as determined by immunohistochemistry (Human Protein Atlas)^45^ (left), and in the adult brain and cerebellum from nine mammals and chicken based on bulk RNA-seq data^46^ (right). The antibodies used for immunohistochemistry were HPA047627 (CPLX4) and HPA011296 (ZP2). Red arrows point to synaptic staining in the granule cell layer. Scale bars, 100 μm. For brain bulk RNA-seq, prefrontal cortex was sampled for primates, and whole brain, except cerebellum, was sampled for non-primates. Dots represent the median, bars indicate the range between the biological replicates. **k**, Expression of genes that were gained or lost in the human or mouse lineage in different cerebellar cell types across development. The expression of the genes that were lost in the human lineage were evaluated in the mouse samples, and *vice versa*. Stages are aligned across species as shown in Fig. 1a; the line indicates the median of biological replicates. **l**, Expression of genes that were gained in the respective cell type in the human or mouse lineage across astroglia subtypes (top) and granule cell states (bottom). The line shows the median of all cells and the dots shows the median of each biological replicate that had at least 50 cells in the respective subgroup. In **e, f**, and **h**, boxes represent the interquartile range, and whiskers extend to extreme values within 1.5 times the interquartile range from the box. diff, differentiating; GC, granule cell; GCP, granule cell progenitor; GL, granule cell layer; GABA DN, GABAergic deep nuclei neurons; glut DN, glutamatergic deep nuclei neurons; PL, Purkinje cell layer; PWM, prospective white matter; RL, rhombic lip; UBC, unipolar brush cell; VZ, ventricular zone.

### SUPPLEMENTARY INFORMATION

#### Supplementary Tables

**Supplementary Table 1**. Samples and libraries.

**Supplementary Table 2**. Clustering and annotation of the mouse dataset.

**Supplementary Table 3**. Clustering and annotation of the human dataset.

**Supplementary Table 4**. Clustering and annotation of the opossum dataset.

**Supplementary Table 5**. Conserved cell state marker genes.

**Supplementary Table 6**. Gene ontology terms associated with the conserved cell state marker genes.

**Supplementary Table 7**. Characterization of orthologous genes for expression trajectories during Purkinje and granule cell differentiation, presence/absence (P/A) expression differences in cerebellar neural cell types across species, and diseases associated with cerebellar functions and cell types.

**Supplementary Table 8**. List of genes that gained expression in human astroglia and are enriched in the rhombic lip and posterior ventricular zone progenitors.

## Notes

https://doi.org/10.11588/data/QDOC4E

### Competing Interest Statement

The authors have declared no competing interest.

## REFERENCES

1. Florio, M. & Huttner, W. B. Neural progenitors, neurogenesis and the evolution of the neocortex. Development 141, 2182–2194 (2014).

2. Rakic, P. Evolution of the neocortex: a perspective from developmental biology. Nat Rev Neurosci 10, 724–735 (2009).

3. Herculano-Houzel, S., Manger, P. R. & Kaas, J. H. Brain scaling in mammalian evolution as a consequence of concerted and mosaic changes in numbers of neurons and average neuronal cell size. Front Neuroanat 8, 77 (2014).

4. Butts, T., Green, M. J. & Wingate, R. J. T. Development of the cerebellum: simple steps to make a “little brain.” Development 141, 4031–4041 (2014).

5. Mariën, P. et al. Consensus paper: Language and the cerebellum: an ongoing enigma. Cerebellum Lond Engl 13, 386–410 (2014).

6. Leto, K. et al. Consensus Paper: Cerebellar Development. Cerebellum 15, 789–828 (2016).

7. Kozareva, V. et al. A transcriptomic atlas of mouse cerebellar cortex comprehensively defines cell types. Nature 598, 214–219 (2021).

8. Kebschull, J. M. et al. Cerebellar nuclei evolved by repeatedly duplicating a conserved cell-type set. Science 370, (2020).

9. Haldipur, P. et al. Spatiotemporal expansion of primary progenitor zones in the developing human cerebellum. Science 366, 454–460 (2019).

10. Carter, R. A. et al. A Single-Cell Transcriptional Atlas of the Developing Murine Cerebellum. Curr Biol 28, 2910–2920.e2 (2018).

11. Vladoiu, M. C. et al. Childhood cerebellar tumours mirror conserved fetal transcriptional programs. Nature 572, 67–73 (2019).

12. Wizeman, J. W., Guo, Q., Wilion, E. M. & Li, J. Y. Specification of diverse cell types during early neurogenesis of the mouse cerebellum. eLife 8, (2019).

13. Aldinger, K. A. et al. Spatial and cell type transcriptional landscape of human cerebellar development. Nat Neurosci 24, 1163–1175 (2021).

14. Welch, J. D. et al. Single-Cell Multi-omic Integration Compares and Contrasts Features of Brain Cell Identity. Cell 177, 1873–1887.e17 (2019).

15. Science, A. I. for B. Allen Developing Mouse Brain Atlas. http://developingmouse.brain-map.org/ (2008).

16. Chemistry, M.-P.-I. of B. GenePaint. https://gp3.mpg.de/ (2004).

17. Osumi-Sutherland, D. et al. Cell type ontologies of the Human Cell Atlas. Nat Cell Biol 23, 1129–1135 (2021).

18. Manno, G. L. et al. Molecular architecture of the developing mouse brain. Nature 596, 92–96 (2021).

19. Cardoso-Moreira, M. et al. Gene expression across mammalian organ development. Nature 571, 505–509 (2019).

20. Millen, K. J., Steshina, E. Y., Iskusnykh, I. Y. & Chizhikov, V. V. Transformation of the cerebellum into more ventral brainstem fates causes cerebellar agenesis in the absence of Ptf1a function. Proc National Acad Sci 111, E1777–E1786 (2014).

21. Hashimoto, R. et al. Origins of oligodendrocytes in the cerebellum, whose development is controlled by the transcription factor, Sox9. Mech Develop 140, 25–40 (2016).

22. Redies, C., Neudert, F. & Lin, J. Cadherins in Cerebellar Development: Translation of Embryonic Patterning into Mature Functional Compartmentalization. Cerebellum 10, 393–408 (2011).

23. Sudarov, A. et al. Ascl1 Genetics Reveals Insights into Cerebellum Local Circuit Assembly. J Neurosci 31, 11055–11069 (2011).

24. Sarropoulos, I. et al. Developmental and evolutionary dynamics of cis-regulatory elements in mouse cerebellar cells. Science 373, eabg4696 (2021).

25. Consalez, G. G., Goldowitz, D., Casoni, F. & Hawkes, R. Origins, Development, and Compartmentation of the Granule Cells of the Cerebellum. Front Neural Circuit 14, 611841 (2021).

26. Parmigiani, E. et al. Heterogeneity and Bipotency of Astroglial-Like Cerebellar Progenitors along the Interneuron and Glial Lineages. J Neurosci 35, 7388–7402 (2015).

27. Cerrato, V. et al. Multiple origins and modularity in the spatiotemporal emergence of cerebellar astrocyte heterogeneity. Plos Biol 16, e2005513 (2018).

28. Bakken, T. E. et al. Comparative cellular analysis of motor cortex in human, marmoset and mouse. Nature 598, 111–119 (2021).

29. Hodge, R. D. et al. Conserved cell types with divergent features in human versus mouse cortex. Nature 573, 61–68 (2019).

30. Tosches, M. A. et al. Evolution of pallium, hippocampus, and cortical cell types revealed by single-cell transcriptomics in reptiles. Science 360, 881–888 (2018).

31. Arendt, D. et al. The origin and evolution of cell types. Nat Rev Genet 17, 744–757 (2016).

32. Schüller, U., Kho, A. T., Zhao, Q., Ma, Q. & Rowitch, D. H. Cerebellar ‘transcriptome’ reveals cell-type and stage-specific expression during postnatal development and tumorigenesis. Mol Cell Neurosci 33, 247–259 (2006).

33. Peng, J. et al. Single-cell transcriptomes reveal molecular specializations of neuronal cell types in the developing cerebellum. J Mol Cell Biol 11, mjy089. (2019).

34. Ross, S. E. et al. Bhlhb5 and Prdm8 Form a Repressor Complex Involved in Neuronal Circuit Assembly. Neuron 73, 292–303 (2012).

35. Britanova, O. et al. Satb2 Is a Postmitotic Determinant for Upper-Layer Neuron Specification in the Neocortex. Neuron 57, 378–392 (2008).

36. Haghverdi, L., Büttner, M., Wolf, F. A., Buettner, F. & Theis, F. J. Diffusion pseudotime robustly reconstructs lineage branching. Nat Methods 13, 845–848 (2016).

37. Butts, T., Wilson, L. & Wingate, R. J. T. Specification of Granule Cells and Purkinje Cells. in Handbook of the Cerebellum and Cerebellar Disorders 89–106 (2013). doi:10.1007/978-94-007-1333-8_6.

38. Bartha, I., Iulio, J. di, Venter, J. C. & Telenti, A. Human gene essentiality. Nat Rev Genet 19, 51–62 (2018).

39. Karczewski, K. J. et al. The mutational constraint spectrum quantified from variation in 141,456 humans. Nature 581, 434–443 (2020).

40. Cacheiro, P. et al. Human and mouse essentiality screens as a resource for disease gene discovery. Nat Commun 11, 655 (2020).

41. Cardoso-Moreira, M. et al. Developmental Gene Expression Differences between Humans and Mammalian Models. Cell Reports 33, 108308 (2020).

42. Stenson, P. D. et al. The Human Gene Mutation Database: towards a comprehensive repository of inherited mutation data for medical research, genetic diagnosis and next-generation sequencing studies. Hum Genet 136, 665–677 (2017).

43. Gröbner, S. N. et al. The landscape of genomic alterations across childhood cancers. Nature 555, 321–327 (2018).

44. Reim, K. et al. Structurally and functionally unique complexins at retinal ribbon synapses. J Cell Biology 169, 669–680 (2005).

45. Uhlén, M. et al. Tissue-based map of the human proteome. Science 347, 1260419–1260419 (2015).

46. Brawand, D. et al. The evolution of gene expression levels in mammalian organs. Nature 478, 343 (2011).

47. Sousa, A. M. et al. Molecular and cellular reorganization of neural circuits in the human lineage. Science 358, 1027–1032 (2017).

48. Northcott, P. A. et al. Subgroup-specific structural variation across 1,000 medulloblastoma genomes. Nature 488, 49–56 (2012).

49. Roussel, M. F. & Stripay, J. L. Modeling pediatric medulloblastoma. Brain Pathol 30, 703–712 (2020).

50. Rueda-Alaña, E. & García-Moreno, F. Time in Neurogenesis: Conservation of the Developmental Formation of the Cerebellar Circuitry. Brain Behav Evol 1–15 (2021) doi:10.1159/000519068.

51. Long, H. K., Prescott, S. L. & Wysocka, J. Ever-Changing Landscapes: Transcriptional Enhancers in Development and Evolution. Cell 167, 1170–1187 (2016).

52. Sarropoulos, I., Marin, R., Cardoso-Moreira, M. & Kaessmann, H. Developmental dynamics of lncRNAs across mammalian organs and species. Nature 571, 510–514 (2019).

53. Paolino, A. et al. Differential timing of a conserved transcriptional network underlies divergent cortical projection routes across mammalian brain evolution. Proc National Acad Sci 117, 10554–10564 (2020).

54. Butts, T., Hanzel, M. & Wingate, R. J. T. Transit amplification in the amniote cerebellum evolved via a heterochronic shift in NeuroD1 expression. Development 141, 2791–2795 (2014).

55. Green, M. J. & Wingate, R. J. Developmental origins of diversity in cerebellar output nuclei. Neural Dev 9, 1 (2014).

56. Palkovits, M. General Neurochemical Techniques. in Microdissection of Individual Brain Nuclei and Areas (eds. Boulton, A. A. & Baker, G. B.) 1–17 (1986). doi:10.1385/0-89603-075-x:1.

57. Krishnaswami, S. et al. Using single nuclei for RNA-seq to capture the transcriptome of postmortem neurons. Nature Protocols 11, 499–524 (2016).

58. Attar, M. et al. A practical solution for preserving single cells for RNA sequencing. Scientific Reports 8, 2151 (2018).

59. Cunningham, F. et al. Ensembl 2019. Nucleic Acids Res 47, gky1113. (2018).

60. Wang, Z.-Y. et al. Transcriptome and translatome co-evolution in mammals. Nature 588, 642–647 (2020).

61. Hu, H. et al. AnimalTFDB 3.0: a comprehensive resource for annotation and prediction of animal transcription factors. Nucleic Acids Res 47, gky822 (2018).

62. Scrucca, L., Fop, M., Murphy, T. B. & Raftery, A. E. mclust 5: Clustering, Classification and Density Estimation Using Gaussian Finite Mixture Models. R J 8, 289–317 (2016).

63. Wolock, S. L., Lopez, R. & Klein, A. M. Scrublet: Computational Identification of Cell Doublets in Single-Cell Transcriptomic Data. Cell Syst 8, 281–291.e9 (2019).

64. Melville, J. uwot: The Uniform Manifold Approximation and Projection (UMAP) Method for Dimensionality Reduction. R package version 0.1.10. https://CRAN.R-project.org/package=uwot (2020).

65. Haghverdi, L., Lun, A. T. L., Morgan, M. D. & Marioni, J. C. Batch effects in single-cell RNA-sequencing data are corrected by matching mutual nearest neighbors. Nat Biotechnol 36, 421–427 (2018).

66. Zeisel, A. et al. Molecular Architecture of the Mouse Nervous System. Cell 174, 999–1014.e22 (2018).

67. Team, S. D. RStan: the R interface to Stan. R package version 2.21.2 http://mc-stan.org/ (2020).

68. Meredith, M. & Kruschke, J. HDInterval: Highest (Posterior) Density Intervals. R package version 0.2.2. https://CRAN.R-project.org/package=HDInterval (2020).

69. Giorgino, T. Computing and Visualizing Dynamic Time Warping Alignments in R : The dtw Package. J Stat Softw 31, (2009).

70. Baglama, J., Reichel, L. & Lewis, B. W. irlba: Fast Truncated Singular Value Decomposition and Principal Components Analysis for Large Dense and Sparse Matrices. R package version 2.3.3. https://CRAN.R-project.org/package=irlba (2019).

71. Young, M. D. & Behjati, S. SoupX removes ambient RNA contamination from droplet-based single-cell RNA sequencing data. Gigascience 9, giaa151 (2020).

72. Korsunsky, I. et al. Fast, sensitive and accurate integration of single-cell data with Harmony. Nat Methods 16, 1289–1296 (2019).

73. Kumar, L. & Futschik, M. E. Mfuzz: A software package for soft clustering of microarray data. Bioinformation 2, 5–7 (2007).

74. Brunson, J. C. & Read, Q. D. ggalluvial: Alluvial Plots in “ggplot2”. R package version 0.12.3. http://corybrunson.github.io/ggalluvial/ (2020).

75. Manno, G. L. et al. RNA velocity of single cells. Nature 560, 494–498 (2018).

76. Yanai, I. et al. Genome-wide midrange transcription profiles reveal expression level relationships in human tissue specification. Bioinformatics 21, 650–659 (2005).

77. Wickham, H. et al. Welcome to the Tidyverse. J Open Source Softw 4, 1686 (2019).

78. Amezquita, R. A. et al. Orchestrating single-cell analysis with Bioconductor. Nat Methods 17, 137–145 (2020).

79. Liu, J. et al. Jointly defining cell types from multiple single-cell datasets using LIGER. Nat Protoc 15, 3632–3662 (2020).

80. Kolde, R. pheatmap: Pretty Heatmaps. https://rdrr.io/cran/pheatmap/ (n.d.).

81. Wickham, H. ggplot2, Elegant Graphics for Data Analysis. (2009) doi:10.1007/978-0-387-98141-3.

82. Wolf, F. A., Angerer, P. & Theis, F. J. SCANPY: large-scale single-cell gene expression data analysis. Genome Biol 19, 15 (2018).

83. Putri, G., Anders, S., Pyl, P., Pimanda, J. & Zanini, F. Analysing high-throughput sequencing data with HTSeq 2.0. https://htseq.readthedocs.io/en/master/ (2021).

